# Aging and Germination of Long-term Stored Seeds: Can MicroRNAs Unlock the Secrets?

**DOI:** 10.1101/2024.07.30.605790

**Authors:** Marta Puchta-Jasińska, Paulina Bolc, Jolanta Groszyk, Maja Boczkowska

## Abstract

**Background:** Small non-coding RNAs appear to be one of the key components of the germination process. To investigate how small non-coding RNAs correlate with germination of seeds with different levels of viability, miRNA-Seq analyses were performed.

**Results:** Our analysis sequencing identified 62 known miRNAs from 11 families and 234 new miRNAs after imbibition process. Among the miRNAs with the highest expression levels, we can mention: miR159, miR168 and miR166. The study placed particular emphasis on miRNAs with significant differences in expression levels at different stages of imbibition and among seeds with different viability. DEG analysis identified 28 miRNAs with significant differences in expression levels, their function was assessed by *in silico* analyses and confirmed by degradome-seq analysis. The expression of miRNAs was verified by qRT-PCR.

**Conclusion:** Our data provides a useful source of information on miRNA during germination long term storage seeds with different viability. The studies suggest that miRNAs are involved in the germination process by their regulation DNA and RNA binding, regulation of developmental process and ribosome.

## Background

Seed germination is a crucial stage in the life cycle of seed plants. It is a transition from the stage of the life cycle that is most resistant to the stage that is most vulnerable to the external environment. Also, it is the point at which a plant becomes a functional component of an ecosystem. In agriculture, seed germination is the basis of crop production. Germination begins when the seed absorbs water (imbibition) and ends when the embryo axis, usually the radicle, emerges through the surrounding structures and is followed by seedling establishment (post germination phase) [1]. Many structural changes and metabolic transformations occur meanwhile and are known as embryo activation [1]. These occur in a specific sequence and in a coordinated manner to ensure the activation of all the basic processes that determine the proper growth and development of the seedling [1]. The initial phase of germination i.e. imbibition occurs spontaneously, driven by the force of the water potential gradient between the dry seed and the surrounding water-rich environment [2]. This phenomenon occurs in both living and dead seeds and is accompanied by massive leakage of cellular solutes because rapid and erratic rehydration can lead to damage to cell membranes and other structures in the seed [3, 4]. Seeds activate numerous repair mechanisms during imbibition to cope with the damage caused by dehydration, storage and, most importantly, rehydration [5]. Repair mechanisms of cell and organelle membranes, proteins, and genomic DNA are activated [6]. The early stage of germination is characterized by a rapid increase in respiration [7]. Primary metabolic pathways are activated, and energy production begins via glycolysis, fermentation, the tricarboxylic acid cycle (TCA), and the oxidative pentose phosphate pathway (OPPP) [8, 9]. Translation of long-lived transcripts stored in dry seeds commences during this phase [10–12]. The entirety of the transcriptional apparatus is stored in dry seeds, with the majority of its components rapidly activated once imbibition occurs [10]. At the end of imbibition, *de novo* mRNA synthesis is also triggered [11]. As water is absorbed by the seed, the gradient of water uptake decreases due to the hydration of cellular components and cell walls. When the rate of water uptake begins to stagnate, seed germination enters phase II, also known as the lag phase [1]. During its course, the main metabolic processes occur in both dormant and nondormant seeds. However, their extent is significantly different. In dormant seeds, cell integrity is restored, mitochondria are repaired, and respiration is initiated with a small breakdown of storage reserves. DNA damage is repaired, and transcription and synthesis of mRNA and biosynthesis of proteins associated with the germination or with dormancy, i.e. encoding transcription factors, hormone metabolism enzymes and signaling proteins, and cell wall modifying enzymes occur [9, 13–16]. In seeds that are not dormant, further processes in preparation for germination take place. These include the completion of repair of DNA damage accumulated during dry storage and the reconstitution of the cytoskeleton associated with the plasma membrane adjacent to the cell wall from tubulin subunits. The synthesis of specific proteins associated with embryo expansion and root protrusion commences, and post-translational modifications of proteins, such as protein phosphorylation and dephosphorylation or ubiquitination, which affect their stability, are initiated [1].

The emergence of the embryo from the surrounding tissues represents the completion of germination in the strict sense of the term. In the majority of species, the radicle is the first structure to emerge subsequent to the physical penetration of surrounding tissues, such as the enclosing endosperm and testa. The third, later phase, also known as the post-germination phase, is the development of the seedling [17]. The process of the radicle emerging from the seed, which is accompanied by a cell elongation, is followed almost immediately by the radicle’s growth, which is due to cell divisions and the elongation of the newly formed cells. Therefore, cell division is a process that occurs subsequent to germination and plays a role in the growth of the axis and the establishment of the seedling [1]. Despite decades of research, many processes involved in seed germination remain insufficiently understood. Therefore, further studies in this area are necessary.

One important piece of this overly intricate puzzle is the fraction of small non-coding RNAs. MicroRNAs (miRNAs) and small interfering RNAs (siRNAs) are two major classes of endogenous small RNAs in plants. They have a significant impact on a multitude of developmental and physiological processes by providing sequence specificity to gene and genome regulation [18]. The mode of action involves silencing genes at both the transcriptional and post-transcriptional levels [19]. The variable levels of sRNAs in plant cells suggest their regulatory role. The best-characterized class of plant sRNAs are miRNAs, which are non-coding, endogenous RNA molecules of 19-24 nucleotides in length [20]. A single miRNA can have multiple target genes, and multiple miRNAs can bind to the same target gene [21, 22]. miRNA can affect translation of mRNAs, promote mRNA cleavage or poly(A) tails shortening [23, 24]. It is estimated that genes encoding miRNAs (*MIRs*) represent about 1% of the predicted genes in higher eukaryotic genomes, and 10-30% of gene expression may be regulated by them [25, 26]. The role of miRNAs in plants includes the regulation of processes responsible for root, leaf, shoot and flower development [27]. In plants, miRNAs are involved in many important seed-related processes, including development, dormancy and germination [28]. Studies of seed germination mechanisms have identified many miRNAs involved in stress responses, phytohormone signaling, antioxidant effects and regulation of key transcription factors, which are thought to be important in seed germination [29, 30]. The precise regulation of miRNAs and their targets remains insufficiently understood, despite their potential significance in the early stages of seed germination.

A high-throughput miRNA analysis of maturating *Brassica napus* seeds revealed the presence of more than 500 conserved miRNAs or variants of unique sequences. The most abundant families were miRNA156, miRNA159, miRNA172, miRNA167, and miRNA158, suggesting their involvement in the network controlling seed development and maturation. A detailed analysis of the results showed that the targets of miRNA173, miRNA400 and miRNA396 are pentatricopeptide repeat-containing proteins (PPRs), suggesting an interaction between RNA-binding proteins and miRNAs during seed germination [31].

Further research on small RNAs will contribute to a more comprehensive understanding of the regulatory network controlled by miRNAs in seed germination [32]. A number of different miRNAs have been identified as regulators of genes encoding activators and repressors of seed germination and seed dormancy [33]. These include miR156, miR158, miR159, miR160, miR164, miR165/166, miR167, miR172, miR395, miR402, and miR417 [34]. Studies by Das et al. suggest that abrupt changes in the expression of several miRNAs, their targets, and interference in signaling between miRNAs and ta-siRNAs (trans-acting siRNAs) contribute to the regulation of seed germination in *Arabidopsis thaliana* L [35]. In maize, the levels of 12 miRNA families were down-regulated during the imbibition phase: miR156, miR159, miR164, miR166, miR167, miR168, miR169, miR172, miR319, miR393, miR394, miR397 [36]. In studies on radish under salinity stress conditions, a negative effect of miR417 on seed germination was observed, while miR395 acts as a regulator of this process. The main role in regulating the dynamic germination process is played by miR159 through modulation of the GA and ABA phytohormone signaling cascades. Overexpression of this molecule has been shown to delay seed germination in radish. It is thought that miR417 affects seed germination through an ABA-dependent pathway, but it is not yet clear whether miR417 regulates germination directly or indirectly by controlling other genes that regulate the response to stressors [36]. It is also worth noting that other phytohormones are regulated by miRNAs during seed germination. For example, nine miRNAs were predicted to be involved in phytohormone signal transduction pathways in barley [37]. Notably, miR156, miR159, miR390, miR164, miR396, and miR319 were predicted to regulate the ethylene pathway [37]. Furthermore, it seems likely that miR172, miR396, and miR319 play a role in cytokinin signaling. In addition, miR393, miR390, miR164, and miR167 appear to be involved in the auxin signaling pathway, whereas miR319 is potentially associated with the jasmonic acid signaling pathway [37].During the germination process, miRNAs also regulate the expression of transcription factors, including ARFs, MYBs, SPLs, NACs and bHLHs [38–46].

Considering the extensive involvement of miRNAs in the regulation of the germination process, the main objective of this study was to determine how reduced viability of naturally aged seeds affects miRNA expression during the first 24 hours of germination.

## Methods

The seed used in the experiment originated from the Botanical Garden - Centre for Biodiversity Conservation of the National Academy of Sciences in Powsin. The analyses were carried out at the National Centre of Plant Genetic Resources, Institute of Plant Breeding and Acclimatization in Radzikow, as part of the Preludium 18 UMO-2019/35/N/NZ9/010446 project funded by the National Science Centre.

The aim of the study was to analyze changes in the microtranscriptome of long-term stored barley seeds with different levels of viability at different stages of germination. The analyses carried out made it possible to correlate the changes in the microtranscriptome with the ability of the grains to initiate the germination process and to complete it correctly.

### Plant material

In this study, barley seeds of the Damazy cultivar were used. A detailed description of all of the plant samples used in this study has been published in a previous study by Puchta et al.[47].

225 seeds were collected from each sample and 25 seeds were placed in 60 mm diameter petri dishes lined with a moist cotton pad for germination. At 6, 12 and 24 hours of germination, embryos with scutellum were collected. Each sample was collected in three independent biological replicates of 25 seeds per replicate.

### miRNA extraction and construction sRNA libraries

The embryonic part from 25 seeds was isolated from the seeds, in order to reduce the amount of starch, which interferes with the isolation of nucleic acids and bulk sample was crushed in a mortar under liquid nitrogen. Isolation of miRNA fractions and construction of miRNAseq libraries were performed as described in detail in Puchta et al. [47].miRNA libraries were sequenced on MiSeq (Illumina) using Reagent Kits v3 (150 cycles) in 51 single-end [48]

### Bioinformatic analysis

The quality of the reads obtained from the sRNA-Seq and degradome-Seq sequencing was assessed using FastQC software [49]. The removal of adapters from the raw reads was performed using the UEA Small RNA Workbench: Adapter Removal software. Subsequently, low-quality reads (Q<30) and reads shorter than 17bp and longer than 25bp were then filtered using The UEA Small RNA Workbench Filter software [50]. The high-quality sequences were then mapped to the *Hordeum vulgare* reference genome (MorexV3_pseudomolecules_assembly), which was obtained from the Ensembl Plants database (Release 57 accessed 17.02.2023) [51]. The known miRNAs and their isoforms were identified through a search of the miRBase database version 22.1 [52]. The UEA Small RNA Workbench miRProf tool was used without mismatch parameter. The UEA Small RNA Workbench miRCat programme was used to identify novel miRNAs [53]. The following parameters were used: (genome hits = 16, hit dist = 200, max gaps = 3, max overlap percentage = 80, max percent unpaired = 50, max unique hits = 3, maxsize = 25, min abundance = 1, min energy = -25, min gc = 20, min hairpin length = 60, min paired = 17, min size = 18, orientation percentage = 80, hairpin extension = 100, p-value = 0.05). If the secondary structure of the sequence met the criteria described by Axtell and Meyers, it was considered a candidate miRNA [54]. The total number of measured values mapped was presented in a normalized to reads per million (RPM). The prediction of the new genes was carried out in silico using a psRNATarget schema V2 software (https://plantgrn.noble.org/psRNATarget/, accessed on 14.03.2023), with expectation score up to 5 and length complementarity was 17 [55].

Quantitative analysis was performed with DESeq2 from the SARTools R package [56]. Dry seed miRNome results from the previous study by Puchta et al. [47]were used as reference. Raw reads were re-mapped to the *H. vulgare* reference genome (MorexV3_pseudomolecules_assembly) to utilize the results. Based on the new mapping, the identification of known and novel miRNAs in dry seeds was performed.

To determine the potential role of *H. vulgare* miRNAs in molecular and biological processes, Gene Ontology (GO) annotations of miRNA target genes were extracted from UniProt identifiers [57]. GO categories represented by *in silico* predicted miRNA targets were visualized using the g:Profiler toolkit [58]. The results are presented in graphical form using the ggplot2 package for the R package [59].

### RT-qPCR quantification

Reverse transcription-qPCR (RT-qPCR) analyses were performed to validate the results obtained by NGS sequencing [60]. Mature miRNA sequence primers were designed using miRprimer software [61] and listed in Supplementary Table 1. MiRNA cDNA synthesis was performed using Mir-X™ miRNA First-Strand Synthesis Kit (Takara Bio, Kusatsu, Japan) according to the manufacturer’s protocol with 400 ng miRNA. The RT-qPCR reaction was performed using the FastStart Essential DNA Green Master Kit (Roche Diagnostics GmbH, Mannheim, Germany) and the LightCycler 96 thermal cycler (Roche, Mannheim, Germany) according to the manufacturer’s protocols. Three biological and three technical replicates were performed for all analyses. Based on the ΔΔCt analysis, the expression of the investigated miRNAs was calculated. Two miRNAs were used as reference, namely miR159, miR166 [47].

## Results

Three sequencing runs on the Illumina MiSeq platform yielded 28,501,642; 29,253,255, and 27,424,301 raw reads, respectively. The raw reads were filtered for quality and length (<17 bp to >25 bp). Filtering resulted in an average of 45% of good quality reads for Rc samples, 39% for Hv samples and 35% for Lv samples. The number of reads matched to r/t RNA was up to 3% for all of reads. On average, 21% of all reads were matched to the genome for the Rc samples, 18% for the Hv samples and 16% for the Lv samples. For details, see Figure 1.

**Figure 1.**
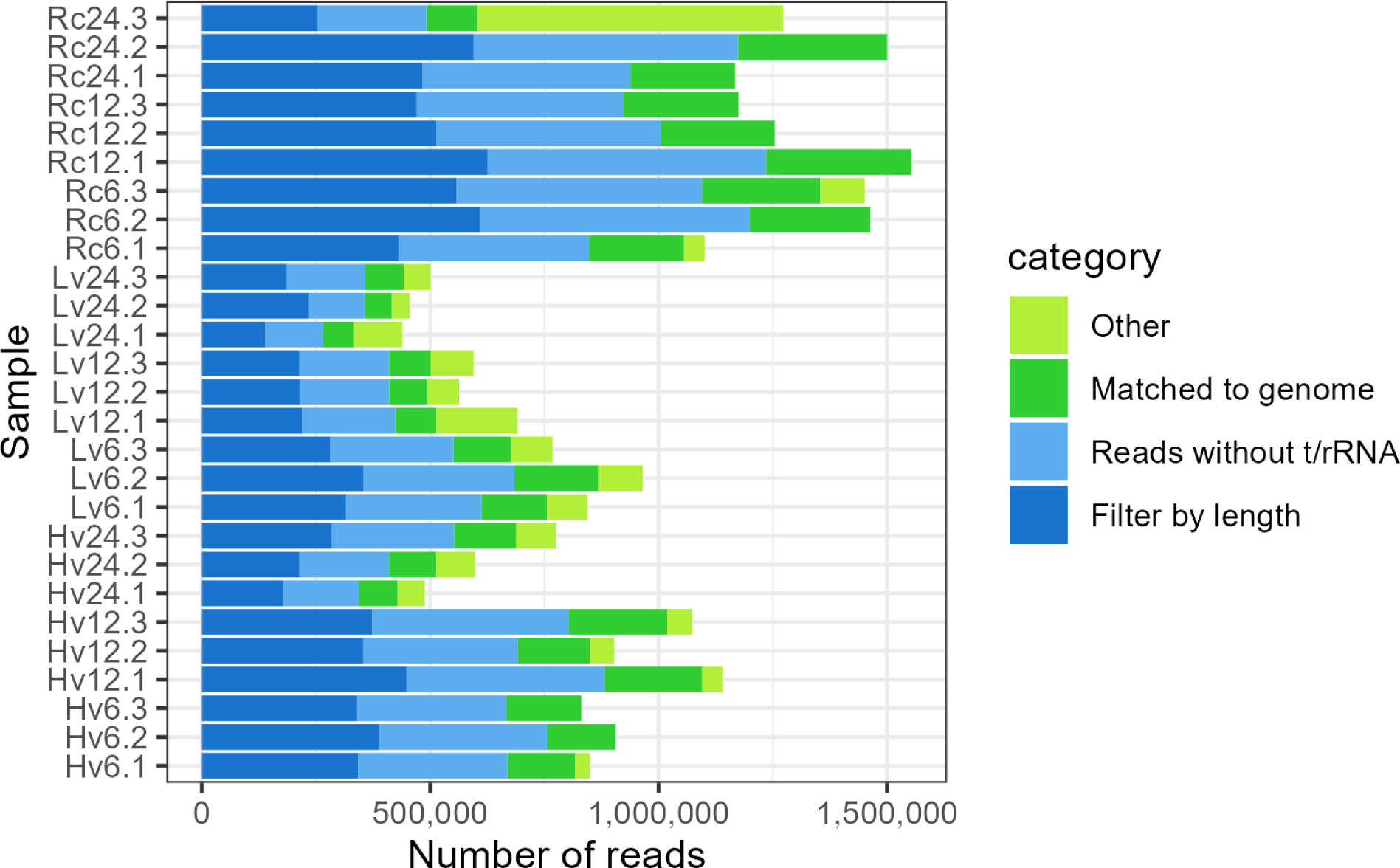
A cumulative bar graph showing the number of raw reads obtained by sRNA-Seq sequencing with a breakdown of reads by group after bioinformatic analysis, including filtering by length, t/rRNA removal, and H. vulgare genome alignment.

The total number of known and new microRNAs detected in the seed samples tested during germination was 296, including 62 known microRNAs and 234 new ones. Of these, 20 known and 13 new miRNAs were detected in all samples. In total in the long-term stored low-viable seed (Lv) sample, 53known and 144 novel miRNAs were observed (Supplementary 2 and 3). Among these, 3 known and 9 novel miRNAs were observed after 6h imbibition, 7 known and 21 novel miRNAs after 12h imbibition, and 2 known and 11 novel miRNAs after 24h imbibition.. Furthermore, 33 known and 56 novel miRNAs were common to all time points examined. In contrast, a total of 172 miRNAs (53 known and 119 novel) were detected at the same time points in samples of long-term stored seeds with high germination capacity (Hv). In detail, 11 known and 28 new unique miRNAs were observed after 6 hours of imbibition, while 3 known and 6 new miRNAs were observed after a further 12 hours, and 3 known and 15 new miRNAs were observed after a further 12 hours (24h in total). The largest number of miRNA occurred in regenerated seed samples (Rc), i.e. there were 58 known and 146 novel miRNA observed in all three investigated time points of germination. Of these, 5 known and 10 novel miRNA were present only in seeds after 6 hours of imbibition, 1 known and 9 novel ones were unique to seed samples after 12 hours of imbibition, and only 3 known and 8 novel ones were present solely after 24 hours of imbibition. For more detailed data, please see Figure 2.

**Figure 2.**
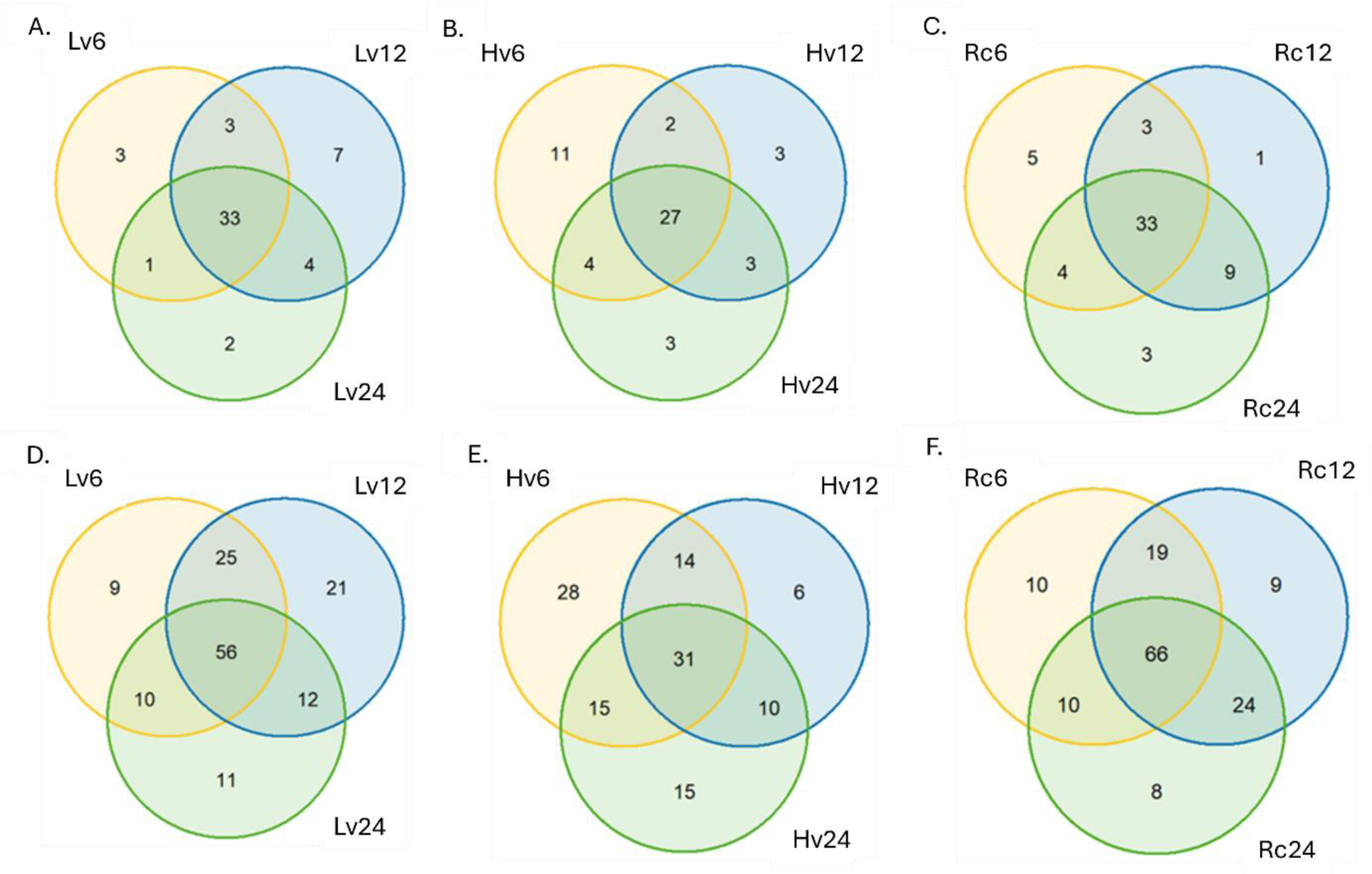
Ven Diagram for number of known and new miRNA detected in barley germination (A. Known miRNA detected in (Hv) highly viable seeds after 6, 12, 24 h germination; B. Known miRNA detected in (Lv) low viable seeds after 6, 12, 24 h germination; C. Known miRNA detected in (Rc) renewed seeds after 6, 12, 24 h germination; D. New miRNA detected in (Hv) highly viable seeds after 6, 12, 24 h germination; E. New miRNA detected in (Lv) low viable seeds after 6, 12, 24 h germination; F. New miRNA detected in (Rc) regenerated seeds after 6, 12, 24 h germination)

In the sequenced sRNA libraries, miRNAs with lengths between 18 and 25 nt were found. Among the known miRNAs, the highest number of miRNAs with a length of 20 nt was observed, that is, 20 known miRNAs, while 18 miRNAs with a length of 19 nt were observed. The lowest number of known miRNAs was observed among the miRNAs with a length of 22 nt (2 ones). There were no known miRNAs longer than 22 nt in the studied material. The highest number of new miRNAs was observed for miRNAs of 21 nt (41 new miRNAs) and 24 nt (29 new miRNAs), while the lowest number of new miRNAs (7) was observed for miRNAs of 25 nt (Figure 3).

**Figure 3.**
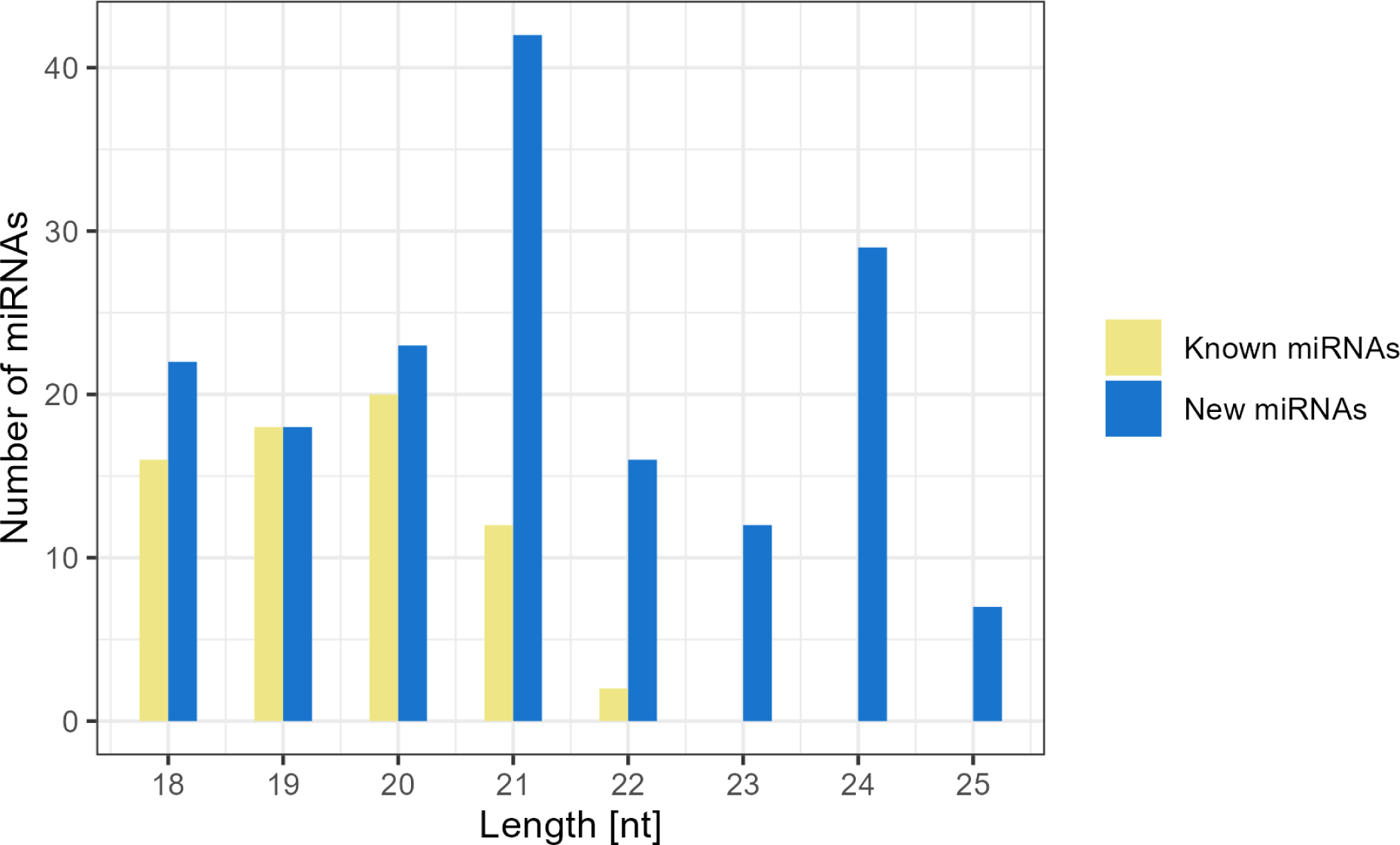
Size distribution of miRNAs in results of next generation sequencing of barley seeds after germination The expression analysis of the identified miRNAs showed that the highest level of expression of both known and novel miRNAs was observed for miRNAs of 21nt length. This represented more than 60% of the total expressed known miRNAs and more than 70% of the total expressed novel miRNAs. The lowest expression was observed for new miRNAs of 23 and 24nt length and 18nt for known miRNAs (Figure 4).

Notably, in the Rc24 sample 21 nt isomiRs represented more than 20 % of both known and novel miRNAs. They were also highly expressed in Lv12, Hv6 and Rc12 samples for both novel and known miRNAs, while representing less than 5% in Hv12 and Hv24 samples. In all samples, less than 5% of known and novel isomiRs of 18, 19 and 20 nt length were present (Supplementary 4A).

The largest number of known isomiRs of 19 nt was observed for the Rc6 sample, which accounts for 14 miRNA members. Among the known miRNAs with a length of 21 nt, the Hv12 sample had the least isomiRs (7 members), while the highest isomiRs in the Lv6, Lv12 and Rc12 and Rc24 samples (12 members). Among the 18 nt isomiRs, more than 10 miRNAs members were observed in Lv12 and all Rc samples. In the analysis of isomiRs in new miRNAs, more than 30 isomiRs of length 21 nt were observed in samples Lv12, Rc12 and Rc24, whereas less than 20 isomiRs were observed in samples Hv12 and Hv24. More than 20 new isomiRs of length 24 were observed in samples Lv12 and Rc12, whereas there were 5 isomiRs in Hv12 and 6 isomiRs in Hv24 sample (Supplementary 4B).

The highest number of isomiRs was observed in the miR168 family, with 12 isomiRs, while 11 isomiRs were identified in the miR5048 family. In the miR159 family, 9 isomiRs were observed, whereas in the 166 family, 8 isomiRs were observed. The fewest isomiRs were identified in the miR6200 (1 isomir) and miR6201 (2 isomiRs) families (Figure 5).

**Figure 4.**
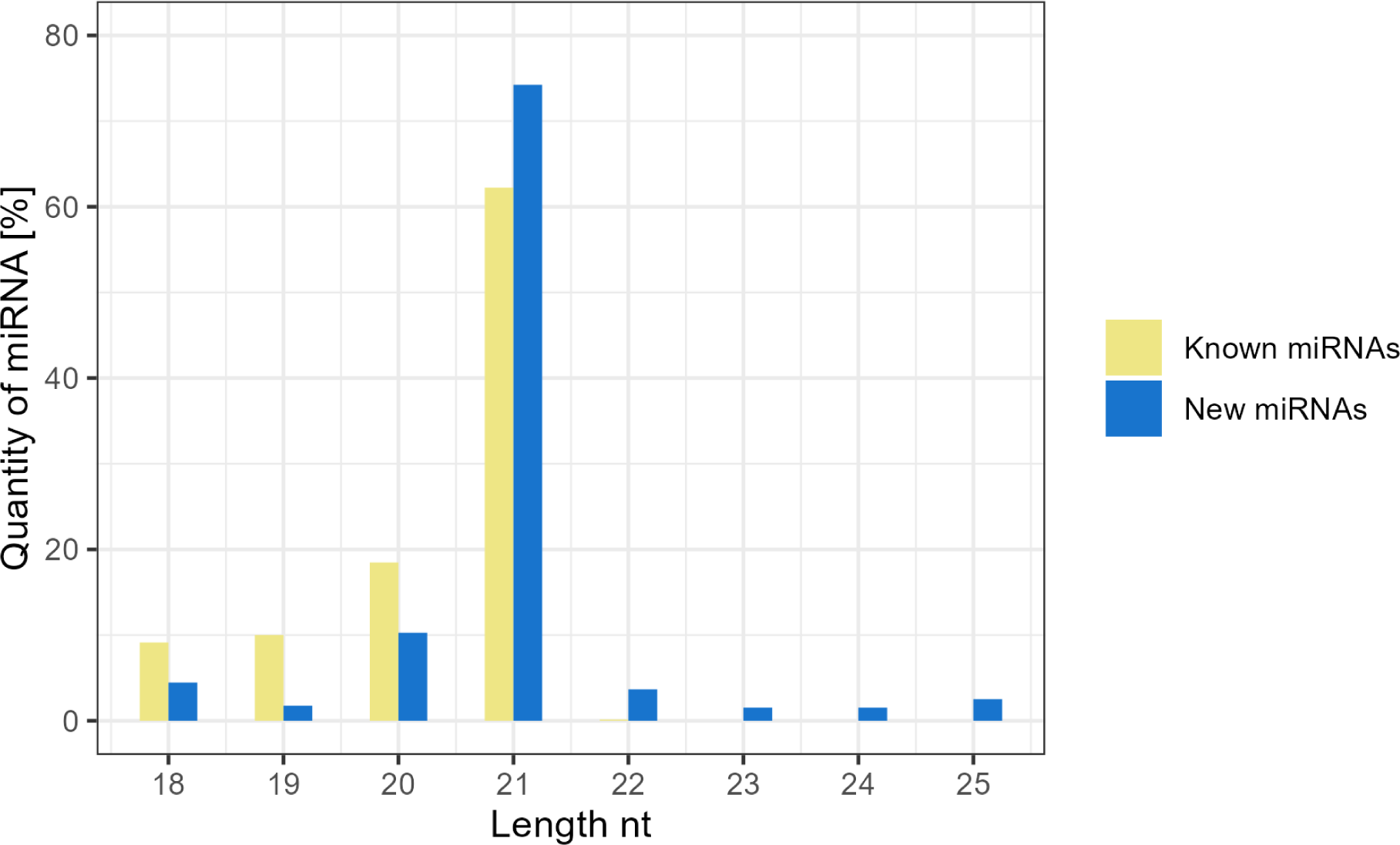
Quantity of known and new miRNA detected in results sRNA sequencing of barley seeds after germination

**Figure 5.**
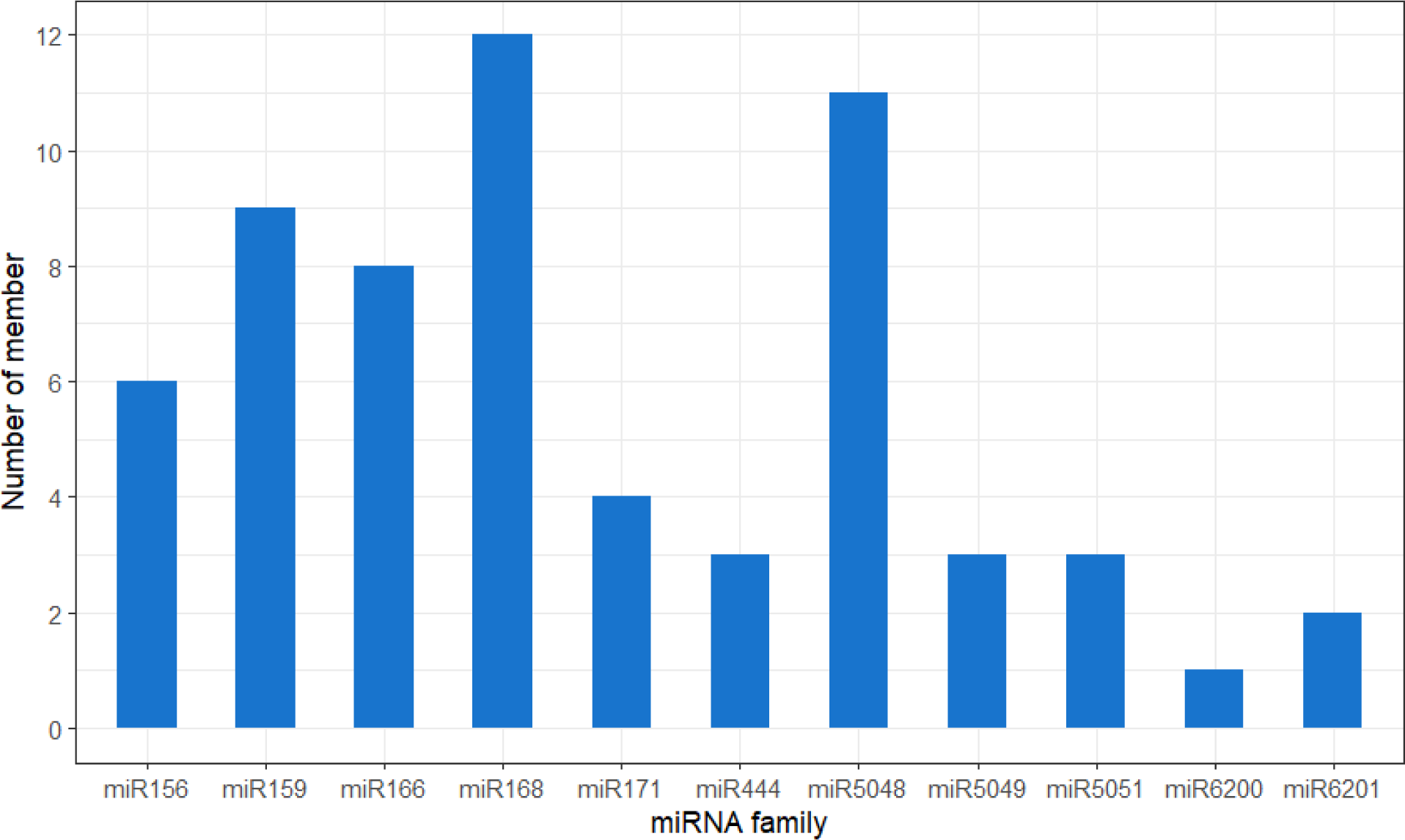
Results of sRNA-Seq analysis of barley seeds during germination showing the number of isomiRs in the miRNA family.

In general, the miR159 family of miR168 and miR166 were found to have the highest expression levels. On average, miR159 was found to be the highest expressed in Rc samples, and in particular in Rc24, it reached 14750.24 RPM. Moreover, its expression level was above 2000.00 RPM in the rest of the samples. It is noteworthy that a record high expression value for miR168 was decimated in the Rc6 sample (45536.00 RPM).

The expression of the miR6200 family was observed only in the Rc12 sample and it was at the level of 1.34 RPM, while miR5049 had the highest expression level in the Rc24 3.47, Rc12 1.62 and Hv6 35431.00. miR444 was the only family with the highest expression in the Lv samples and the lowest in the Rc ones. See Figure 6 for more details.

**Figure 6.**
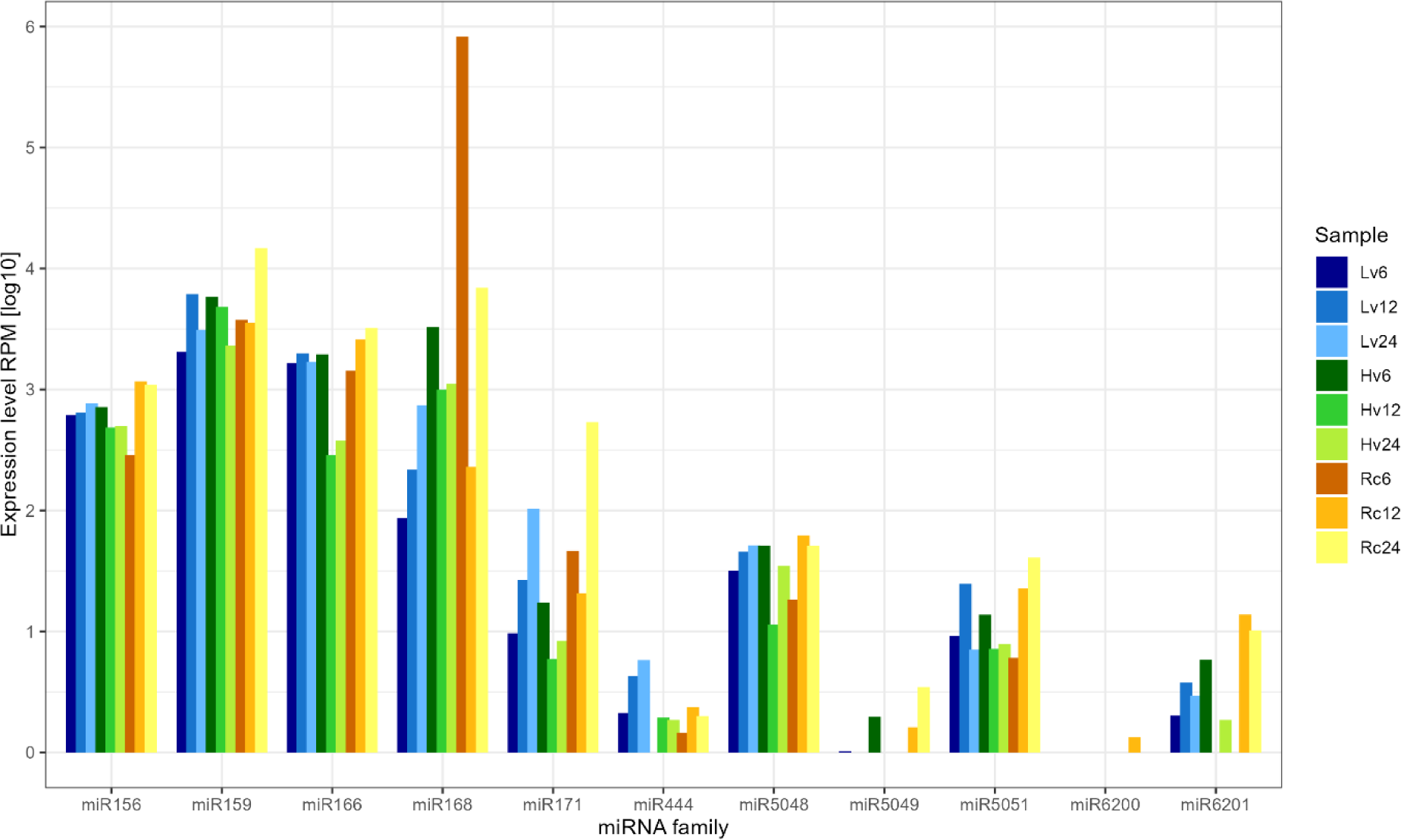
Expression level of miRNAs obtained by NGS sequencing during germination after 6, 12 and 24 h imbibition. Rc-renewed seed sample; Lv-low viable seed; Hv- highly viable seed

Regardless of germination time and sample viability, families with constant expression levels were observed. For individual isomiRs, higher differences in expression levels between samples were observed (Figure 7). The highest expression of miR156_a was found in the Rc6 (43466.00 RPM), whereas in the other samples the expression was close to zero. miR159_b showed high expression only in the Hv6 (45444.00 RPM), whereas miR159_d showed high expression in the Lv6 (29646.00 RPM), Hv12 (35462.00 RPM), Rc6 (16862.00 RPM) and Rc12 (45445.00 RPM). The presence of miR166_c with an expression of more than 25000.00 RPM was found in the Hv6, Hv24 and Rc12, whereas miR166_e showed a high expression only in the Lv24 (45.00 RPM). IsomiR444_c was only expressed in the Hv6 (45444.00 RPM), miR548_d in the Hv12 (19664.00 RPM) and miR5048_c in the Lv24 (32143.00 RPM).

**Figure 7.**
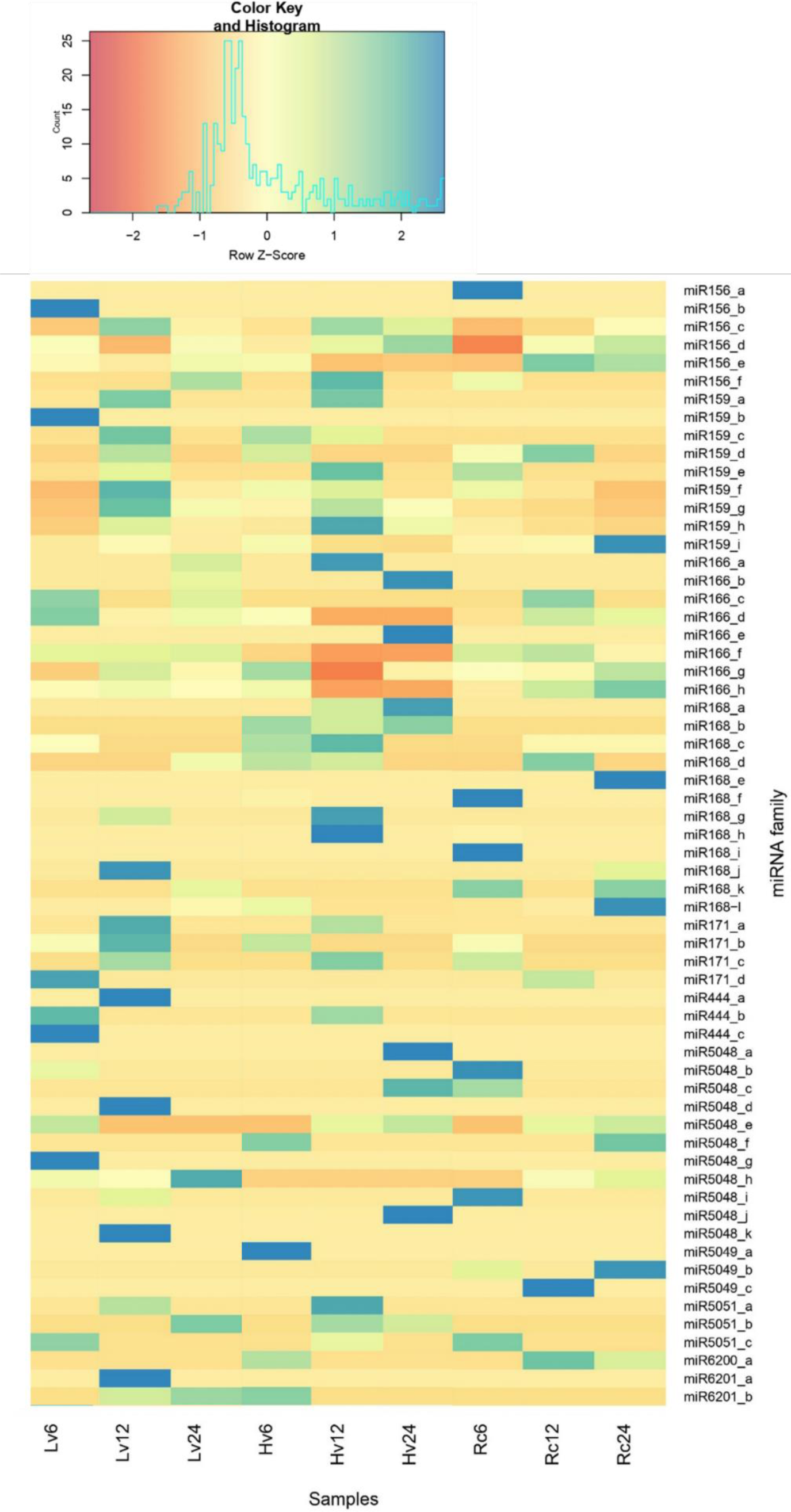
The heatmap for known miRNA expression during 6, 12, 24h imbibition

The evaluation of the differences in the expression levels of the new miRNAs focused on isomiRs with expression levels above 1 (Figure 8). The newly identified miR-10H showed expression only in Hv6 (1.19 RPM) and Lv6 (2.82 RPM) samples. Expression of miR-19H was only observed in Hv samples (8.91 RPM - Hv6; 5.80 RPM - Hv12 and 1.20 RPM - Hv24). miR-154H was expressed at levels above 50.00 RPM, with the exception of samples Hv12 and Hv24, where expression was found at around 3.00 RPM. The presence of miR-160H was observed in Hv and Lv samples after 6 hours of imbibition, where expression was 14.67 and 11.70 RPM, in Lv12 and Rc12 (6.08 and 35.27 RPM) and in Lv24 and Rc24 (10.28 and 31.41 RPM). miR-201H was only expressed in Hv12 (4.11 RPM) and Hv24 (2.54).

**Figure 8.**
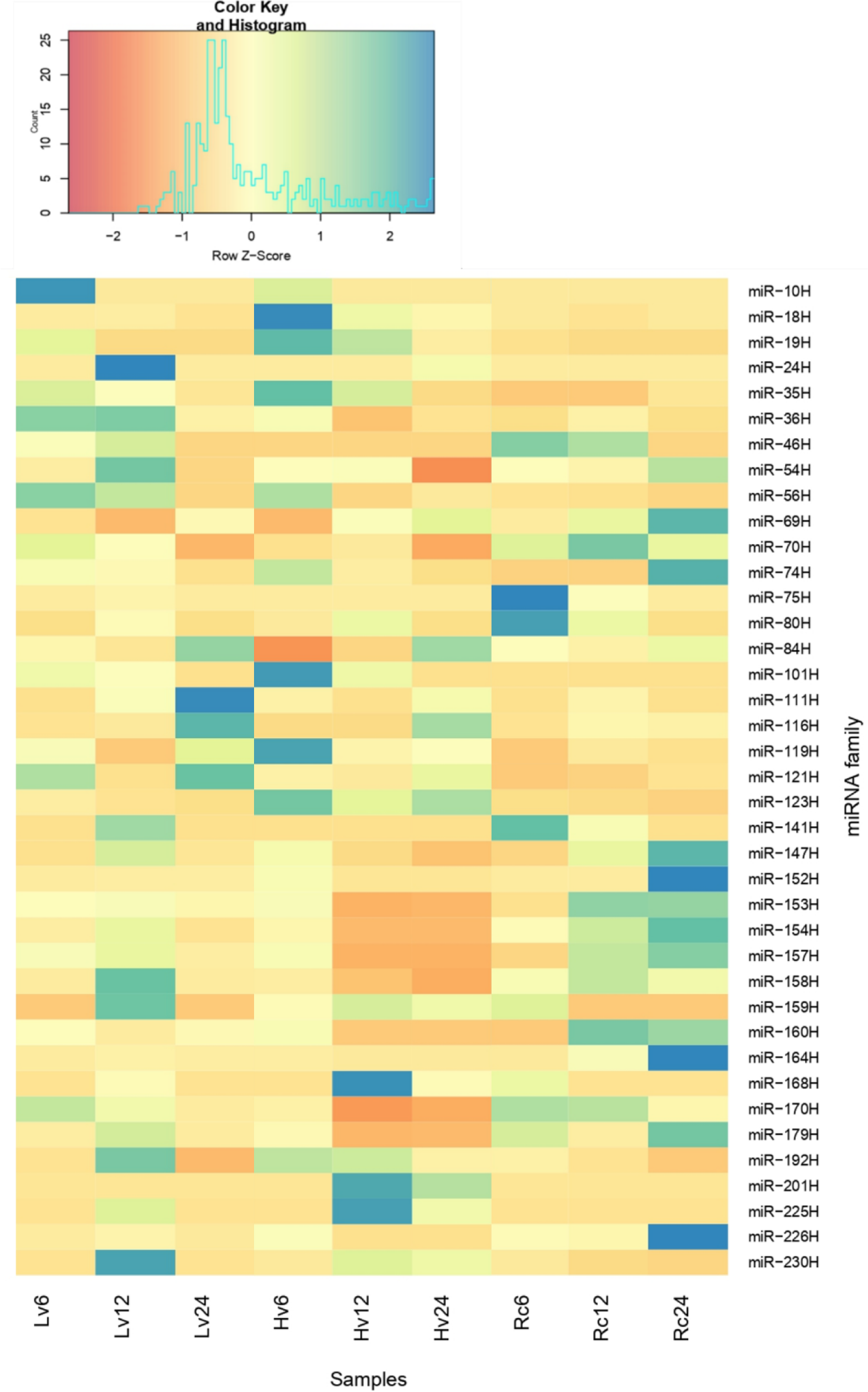
The heatmap for selected new miRNA expression above 1 during 6, 12, 24h imbibition

Differential expression (DE) analysis was performed for comparison of miRNA levels in barley seeds with different viability during 24 h of imbibition. Based on the results, 10 miRNAs, including six novel miRNAs and four known miRNAs, were selected for verification by RT-qPCR analysis. In the NGS and RT-qPCR reactions, a similar expression profile was observed for most of the validated miRNAs. miR-191H and miR-225H were amplified in Hv12 and Hv24 samples and miR-56H in Hv6 and Hv24. miR159g, miR166e, miR166f and miR156c were amplified in all samples tested at all time poins. miR-10H was amplified only in Lv and Hv samples at 6 hours of imbibition (Fig. 9).

**Figure 9.**
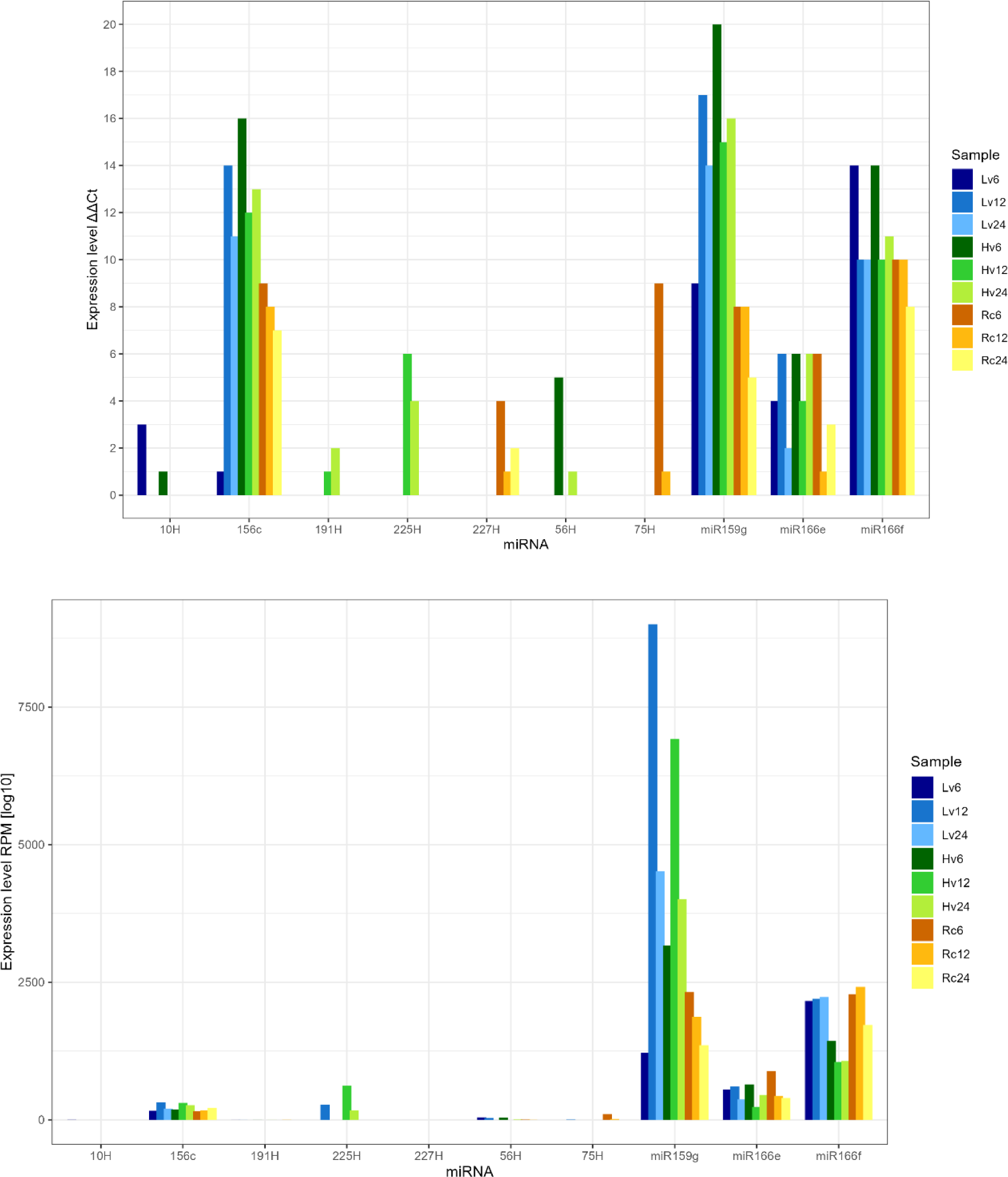
Validation of the expression level of a selected miRNA in barley after 6, 12 and 24 hours of imbibition. Rc-renewed seeds sample; Lv- low viable seeds; Hv- highly viable seeds; **A.** performed by RT-qPCR, **B.** performed by NGS sequencing.

**Figure 9.**
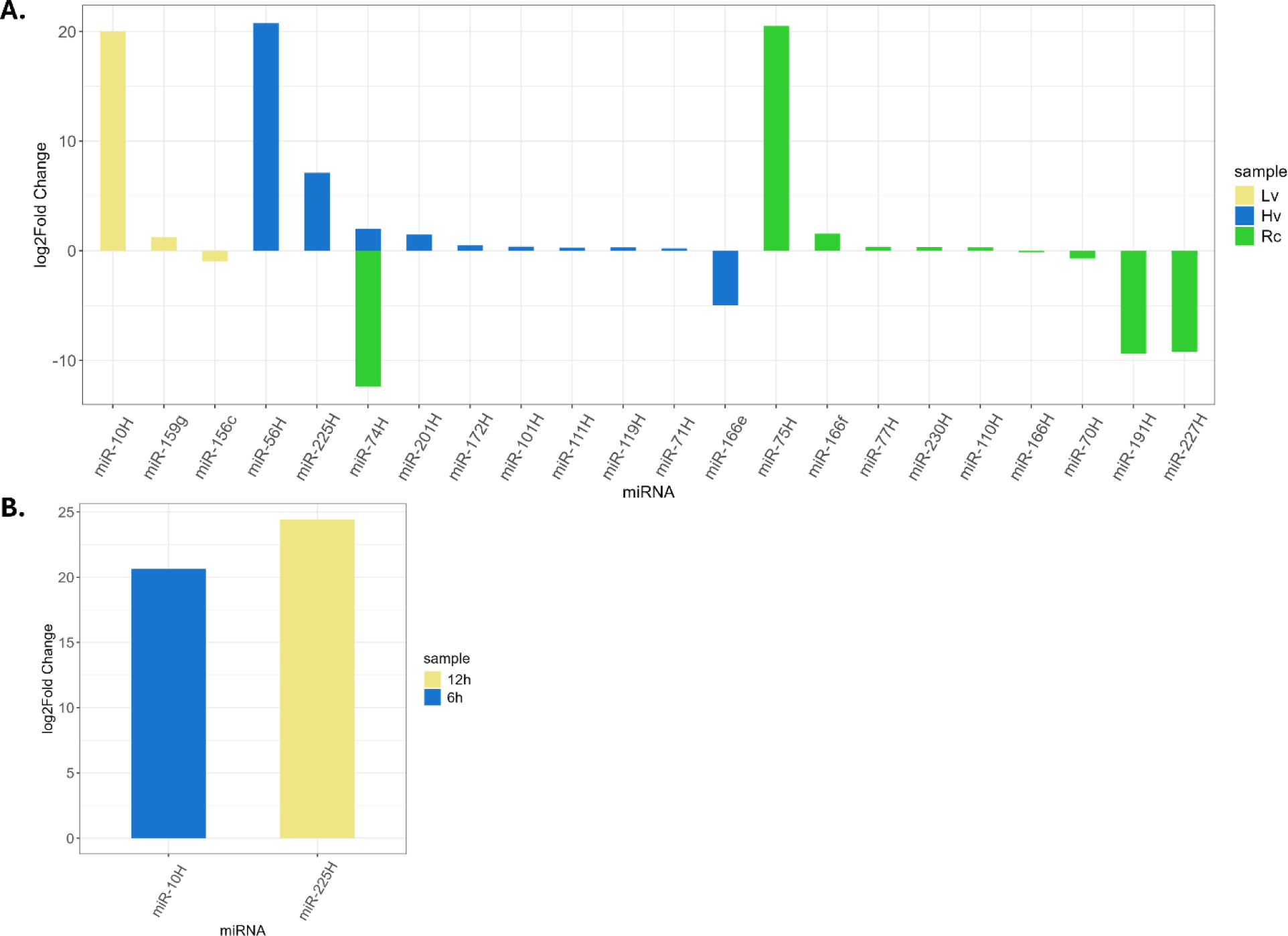
miRNA with significant changes in level expression after DEG analysis (**A**. Hv - miRNA with DEG in seeds with high viability after 6,12,24 h imbibition; Lv - miRNA with DEG in seeds with low viability after 6,12,24 h imbibition, Rc- miRNA with DEG in renewed seeds after 6,12,24 h imbibition, **B**. 12h- miRNA with DEG in Rc, Hv and Lv sample after 12 h imbibition; 6h- miRNA with DEG in Rc, Hv and Lv sample after 6h imbibition).

A differential expression gene (DEG) analysis was performed on both known and newly identified miRNAs. Two experimental designs were examined: changes occurring between samples of different viability at a specific moment of imbibition, and for each sample in successive stages of imbibition. A total of 10 miRNAs with significantly different expression levels at different imbibition stages in Rc samples were identified; five were up-regulated while five were down-regulated (Figure 9). The up-regulated miRNAs with the highest log2 fold-change were miR-75H (20.5; Rc), miR-10H (20.1 Lv) and miR-56H (20,8, Hv). The down-regulated miRNAs with the highest log2 fold-change were miR-74 (-12.4, Rc), miR-191H (-9,4, Rc) and miR-227H (-9,2, Rc). In the Hv samples at different stages of imbibition, the log2 fold-change value was the highest for miR-56H (20.8) and the lowest for miR166e (-4.9). Furthermore, three miRNAs with statistically significant differences in expression levels at different stages of imbibition were identified in the Lv sample. In addition, significant changes in expression levels at the specific time of imbibition were found for miR-10H and miR-225H, which were up-regulated after 6 and 12 hours of imbibition (log2 fold-change 20.6 and 24.4, respectively) (Figure 9).

*In silico* miRNA targets were used to identify miRNA-associated proteins and functions with significantly different expression levels, as shown in Table 1. Functional analysis revealed three potential targets for miR-10H at 6 hours and two for miR-225H at 12 hours of imbibition that differentiating seed samples with different viability levels at specific time points. In total, the analysis identified 8 potential targets for 5 miRNAs in the Rc, 6 potential targets for 3 miRNAs in the Lv, and as many as 14 potential targets for 8 miRNAs in the Hv sample. Notably, the Clu domain-containing protein (A0A8I6Y4N0) transcript was targeted by three different miRNAs, miR-159g in Lv, miR-225H in Hv and miR-227H in Rc. The similarity of the sequences of the last two miRNAs suggests that they may be isomiRs. The MD-2-related lipid-recognition domain-containing protein (A0A8I7B5W4) transcript was identified as a potential target for two miR-227H (Hv) and miR-191H (Hv and Rc). In contrast, two potential targets were identified for miR-75H (Rc). Both were transcripts of genes encoding MDR-like ABC transporter proteins (A0A8I6X084 and A0A8I7B4F3). Four potential targets were identified for miR156c in Lv. All were transcripts of genes encoding the SBP-type domain-containing proteins (A0A8I6XRG1, UPI00162D8BA4, A0A8I6WHB1 and A0A8I7BBH7).

**Table 1.** Target prediction from miRNA with significant level of expression and their encoding proteins.

| miRNA | log2FC | p.adj | Target Accession | Expectation | Protein ID | Protein |
| --- | --- | --- | --- | --- | --- | --- |
| Lv |  |  |  |  |  |  |
| miR159g | 1,229 | 0,034894161 | HORVU.MOREX.r2.6HG0505470.1 | 1 | A0A8I6Y4N0 | Clu domain-containing protein |
| miR-10H | 20,006 | 4,05149E-06 | HORVU.MOREX.r2.5HG0429630.1 | 3 | A0A8I6XX43 | Trafficking protein particle complex subunit 8 |
| miR156c | -0,96 | 0,014083698 | HORVU.MOREX.r2.5HG0406890.1 | 1 | A0A8I6XRG1 | SBP-type domain-containing protein |
|  |  |  | HORVU.MOREX.r2.7HG0564420.1 | 1 | UPI00162D8BA4 | squamosa promoter-binding-like protein 17 isoform X1 |
|  |  |  | HORVU.MOREX.r2.2HG0157010.1 | 1 | A0A8I6WHB1 | SBP-type domain-containing protein |
|  |  |  | HORVU.MOREX.r2.6HG0473260.1 | 1 | A0A8I7BBH7 | SBP-type domain-containing protein |
| Hv |  |  |  |  |  |  |
| miR-56H | 20,76 | 1,91E-08 | HORVU.MOREX.r2.7HG0540020.1 | 1 | A0A8I6Z190 | GDSL esterase/lipase |
| miR-225H | 7,118 | 0,002109966 | HORVU.MOREX.r2.6HG0505470.1 | 1,5 | A0A8I6Y4N0 | Clu domain-containing protein |
|  |  |  | HORVU.MOREX.r2.1HG0024800.1 | 1,5 | A0A287F185 | Uncharacterized protein |
| miR-74H | 1,995 | 0,012285495 | HORVU.MOREX.r2.5HG0405920.1 | 2 | M0X912 | AAA+ ATPase domain-containing protein |
|  |  |  | HORVU.MOREX.r2.3HG0232580.1 | 2 | A0A8I6XF02 | Matrin-type domain-containing protein |
| miR-201H | 1,473 | 0,015042169 | HORVU.MOREX.r2.1HG0065170.1 | 1 | Q70YI6 | phospholipase D |
|  |  |  | HORVU.MOREX.r2.1HG0040430.1 | 1 | A0A8I6WG72 | phospholipase D |
|  |  |  | HORVU.MOREX.r2.4HG0284980.1 | 1 | A0A8I6XX46 | TF-B3 domain-containing protein |
| miR-101H | 0,376 | 0,034548527 | HORVU.MOREX.r2.5HG0389040.1 | 2 | B3W6G6 | Putative transcriptional activator DEMETER |
|  |  |  | HORVU.MOREX.r2.6HG0505760.1 | 2 | A0A8I7BHC5 | PHD-type domain-containing protein |
| miR-119H | 0,305 | 0,035977764 | HORVU.MOREX.r2.2HG0153460.1 | 2 | A0A8I6WX50 | Cationic amino acid transporter C-terminal domain-containing protein |
| miR-227H | -0,38 | 0,035977764 | HORVU.MOREX.r2.6HG0505470.1 | 1,5 | A0A8I6Y4N0 | Clu domain-containing protein |
|  |  |  | HORVU.MOREX.r2.3HG0201760.1 | 1,5 | A0A8I7B5W4 | MD-2-related lipid-recognition domain-containing protein |
| miR-191H | -0,298 | 0,041916293 | HORVU.MOREX.r2.2HG0125310.1 | 1 | A0A8I7B5W4 | MD-2-related lipid-recognition domain-containing protein |
| Rc |  |  |  |  |  |  |
| miR-191H | -9,397 | 5,01E-10 | HORVU.MOREX.r2.2HG0125310.1 | 1 | A0A8I7B5W4 | MD-2-related lipid-recognition domain-containing protein |
| miR-227H | -9,218 | 2,75E-09 | HORVU.MOREX.r2.6HG0505470.1 | 1,5 | A0A8I6Y4N0 | Clu domain-containing protein |
|  |  |  | HORVU.MOREX.r2.1HG0024800.1 | 1,5 | F2EB30 | YTH domain-containing family protein |
| miR-75H | 20,505 | 3,34E-09 | HORVU.MOREX.r2.5HG0404560.1 | 2 | A0A8I6XTH8 | SAM-dependent MTase RsmB/NOP-type domain-containing protein |
|  |  |  | HORVU.MOREX.r2.2HG0079610.1 | 2 | A0A8I6X084 | MDR-like ABC transporter |
|  |  |  | HORVU.MOREX.r2.2HG0079600.1 | 2 | A0A8I7B4F3 | MDR-like ABC transporter |
| miR-77H | 0,368 | 0,031983925 | HORVU.MOREX.r2.6HG0474040.1 | 1,5 | A0A287TTV1 | Inositol-1-monophosphatase |
| miR-230H | 0,335 | 0,034150503 | HORVU.MOREX.r2.7HG0566090.1 | 1,5 | A0A8I6YC91 | DUF632 domain-containing protein |
| Changes in group (Lv, Hv and Rc) after 6, 12 and 24 h imbibition |  |  |  |  |  |  |
| 6 h |  |  |  |  |  |  |
| miR-10H | 20,632 | 1,08E-06 | HORVU.MOREX.r2.5HG0429630.1 | 2 | F2CX01 | Protoporphyrinogen oxidase |
|  |  |  | HORVU.MOREX.r2.7HG0602590.1 | 2 | A0A8I6Z793 | Putative tRNA (cytidine(32)/guanosine(34)-2'-O)-methyltransferase |
|  |  |  | HORVU.MOREX.r2.5HG0374850.1 | 2 | A0A8I6YEI0 | Protein SUPPRESSOR OF GENE SILENCING 3 |
| 12h |  |  |  |  |  |  |
| miR-225H | 24,42 | 2,70E-10 | HORVU.MOREX.r2.6HG0505470.1 | 1,5 | A0A8I6Y4N0 | Clu domain-containing protein |
|  |  |  | HORVU.MOREX.r2.1HG0024800.1 | 1,5 | F2EB30 | YTH domain-containing family protein |

Analysis of GO annotation of miRNA targets with significant differences in expression levels for highly viable (Hv) seeds at different imbibition times showed that among the genes involved in molecular functions, more than 25% were responsible for GO:0003677 DNA binding and GO:0003700 DNA-binding transcription factor activity and their activity localization was mainly in the nucleus. For miRNA target genes in low viability (Lv) seeds, metal ion binding GO:0046872 and DNA binding GO:0003677 were predominant among molecular functions (more than 35% each). The activity was localized to the nucleus in 20% of the targets. Among the miRNAs with significant differences in expression levels at different stages of imbibition in regenerated samples (Rc), the molecular functions of the targets were mainly lipid binding GO:0008289 (over 35%), DNA-binding transcription factor activity GO:0003700 (over 30%), ATP binding GO:0005524 (over 25%) and DNA binding GO:0003677. As before, the targets were localized in the nucleus GO:0005634 (45%) (Fig. 10).

**Figure 10.**
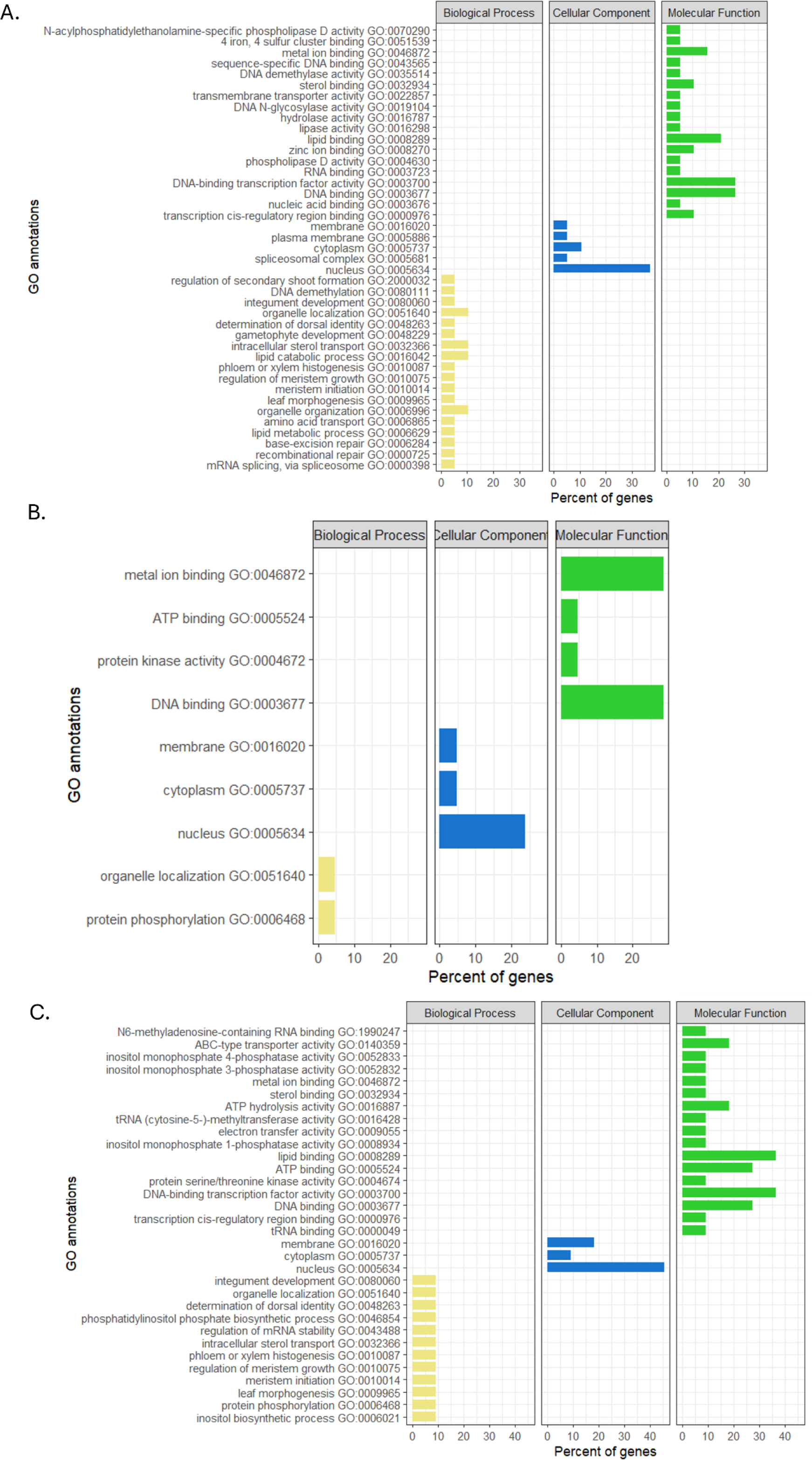
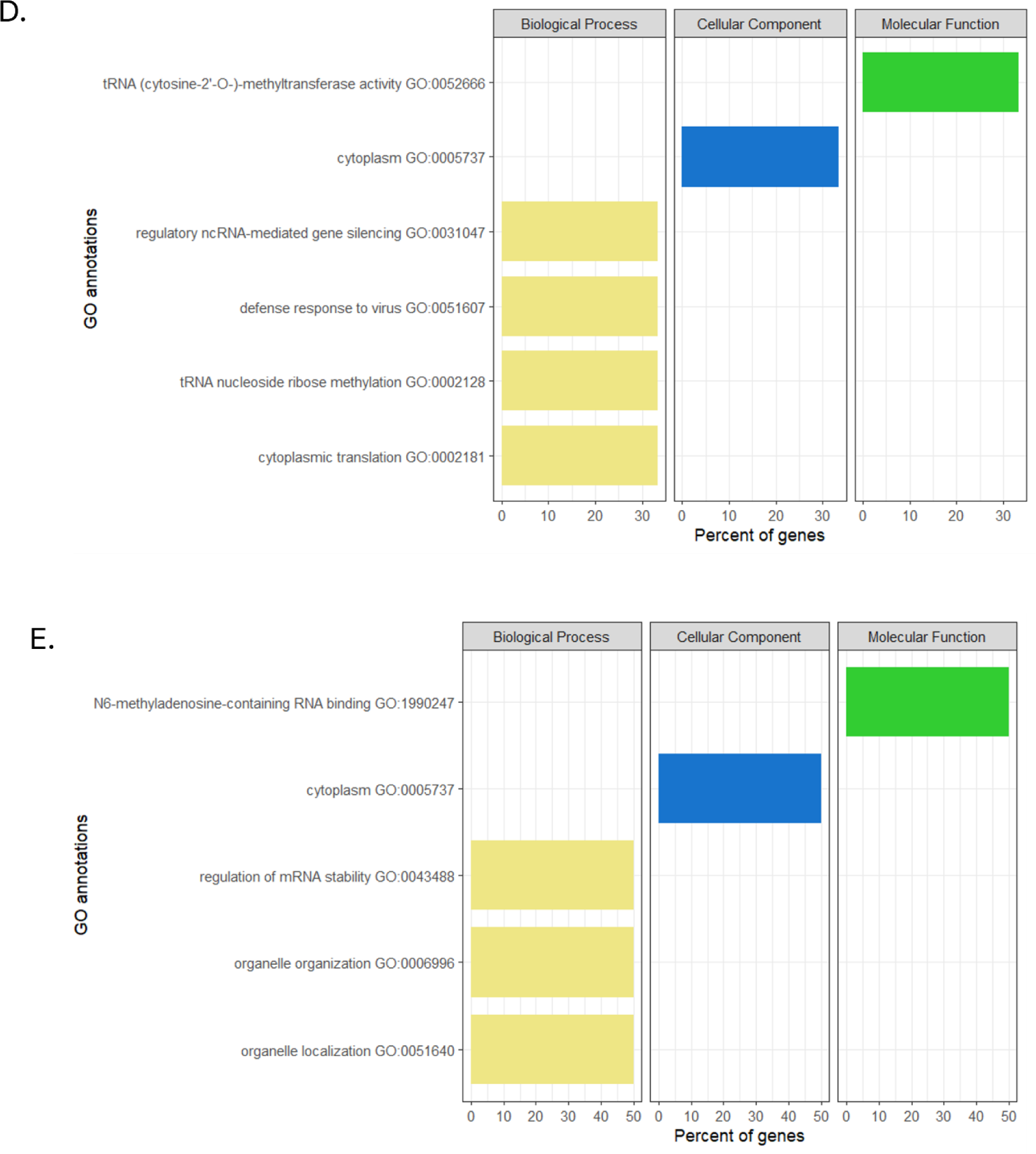
GO enrichment analysis, targets for miRNA with significant expression level. GO analysis classified into three group: molecular function, biological process and cellular component. (A. Hv - miRNA with DEG in seeds with high viability after 6, 12, 24 h imbibition; B. Lv - miRNA with DEG in seeds with low viability after 6, 12, 24 h imbibition, C. Rc- miRNA with DEG in renewed seeds after 6, 12, 24 h imbibition, D. 6h - miRNA with DEG in Rc, Hv and Lv sample after 12 h imbibition; E. 12h- miRNA with DEG in Rc, Hv and Lv sample after 6h imbibition).

Function analysis of miRNAs target genes that showed significant differences in expression levels during 6 hours of imbibition compared to dry seeds showed that over 30% of these genes were related to cytoplasm GO:0005737 and molecular function, such as the tRNA (cytosine 2’-O-) methyltransferase activity, cytoplasmic translation GO:0002181, and tRNA nucleoside-ribose methylation GO:0002128. In contrast, after 12 h of imbibition, cell localization was still associated with the cytoplasm (more than 50%), but their molecular function was related to N-6-methyladenosine-containing RNA binding GO:1990247 and regulation of mRNA stability GO:0043488 (Fig. 10).

GO analysis of potential miRNA targets showed that the majority of up-regulated target genes were involved in for lipid binding processes (p adj. 1.378×10-2). Up-regulated miRNAs were also associated with functions related to developmental regulation and DNA-binding transcription factor activity. A statistically significant down-regulated miRNA was associated with target genes involved in DNA binding (p.adj. 8.958×10-8). The association of target genes responsible for the binding of organic cyclic compounds and the binding of heterocyclic compounds with nucleic acid binding, which is related to the molecular function of DNA binding, was observed. The nuclear localized cellular processes were associated with target genes related to membrane-bound organelles and intercellular membrane-bound organelles. The target genes associated with the up-regulated miRNA were associated with the activity of DNA-binding transcription factors.

**Figure 11.**
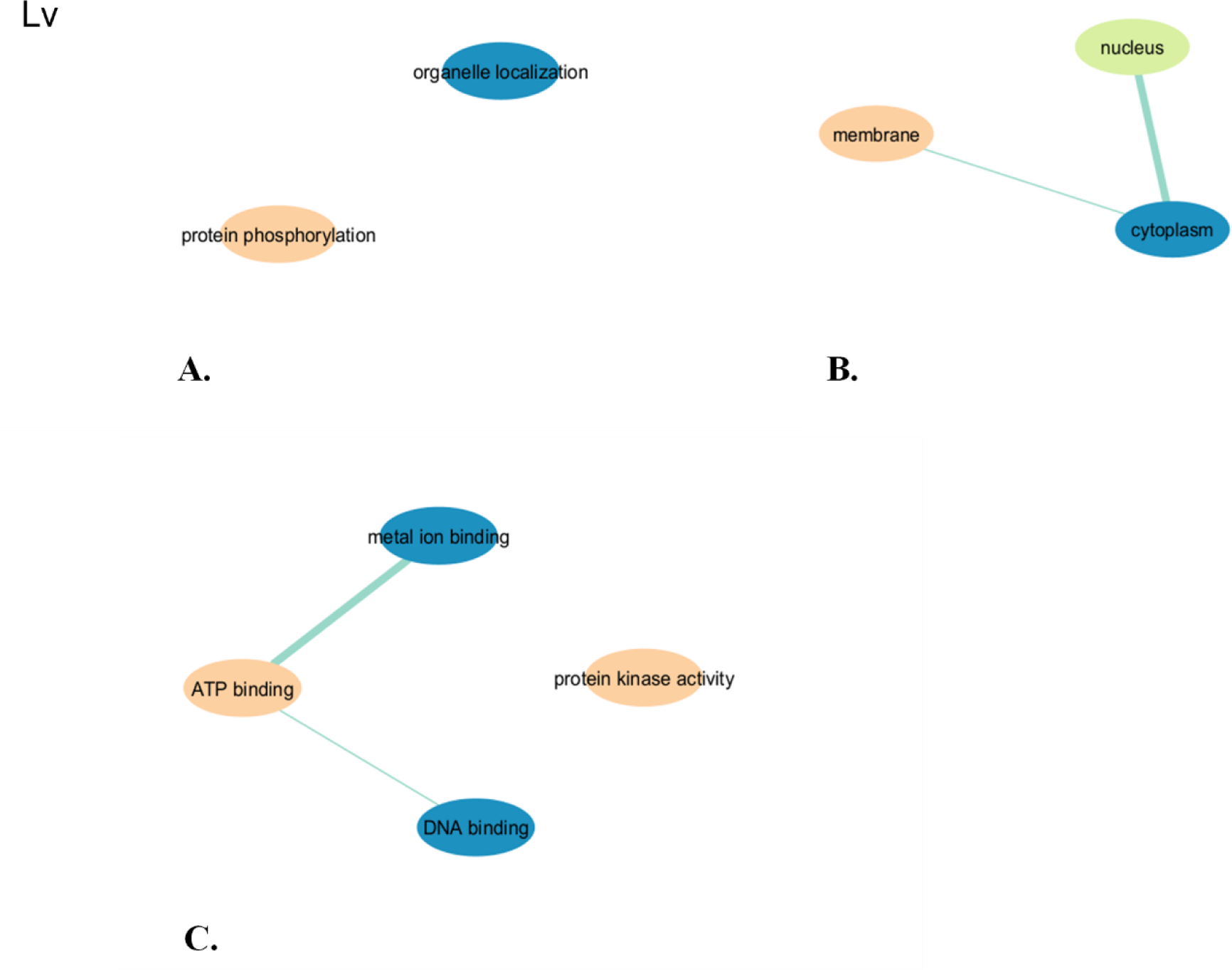
GO analysis of potential miRNA targets of the highly viable seeds sample (Lv) for biological, cellular and molecular functions: **A.** biological functions, **B.** cellular functions, **C.** molecular functions.

**Figure 12.**
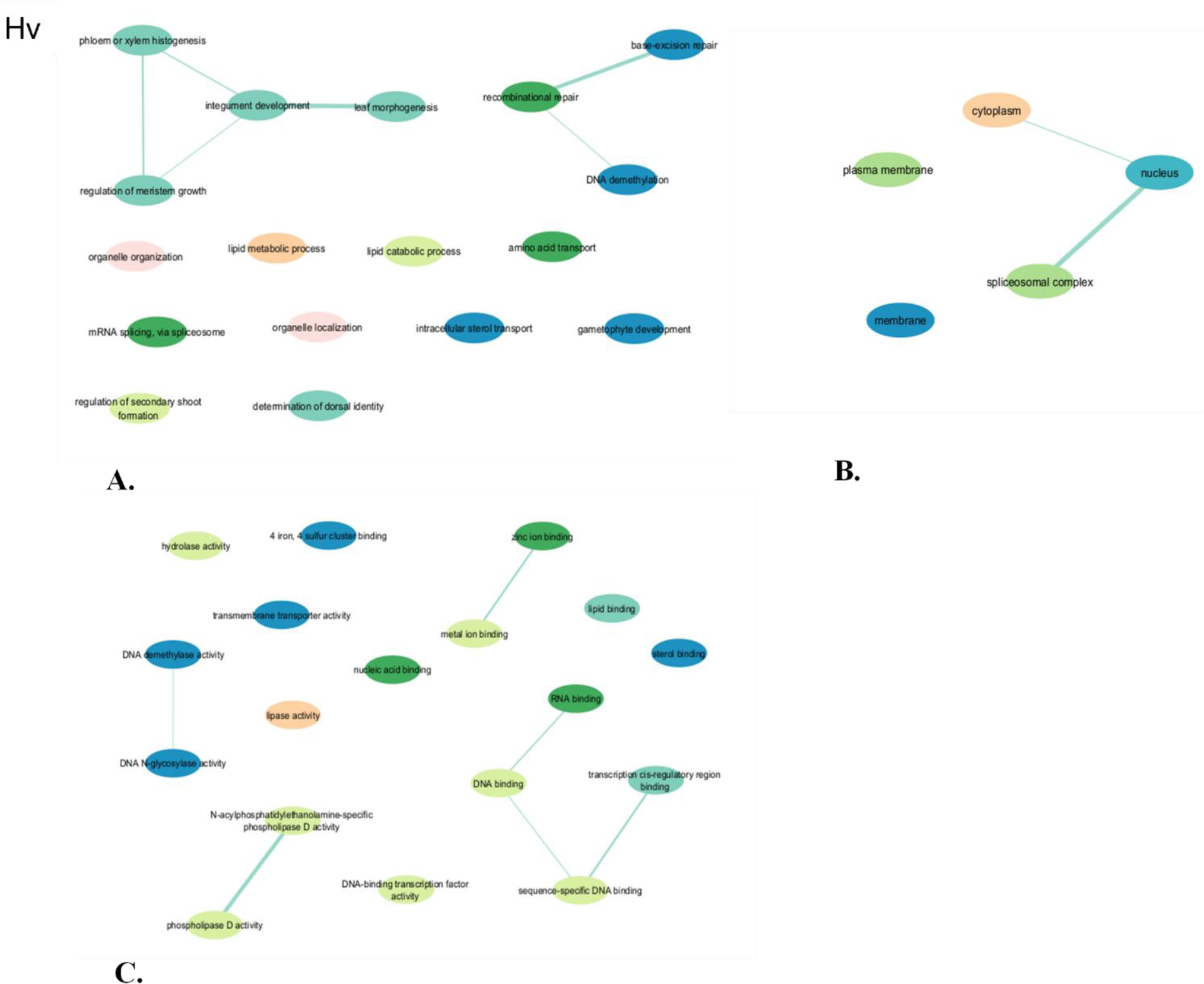
GO analysis of potential miRNA targets of the low viable seeds sample (Hv) for biological, cellular and molecular functions: **A.** biological functions, **B.** cellular functions, **C.** molecular functions.

**Figure 13.**
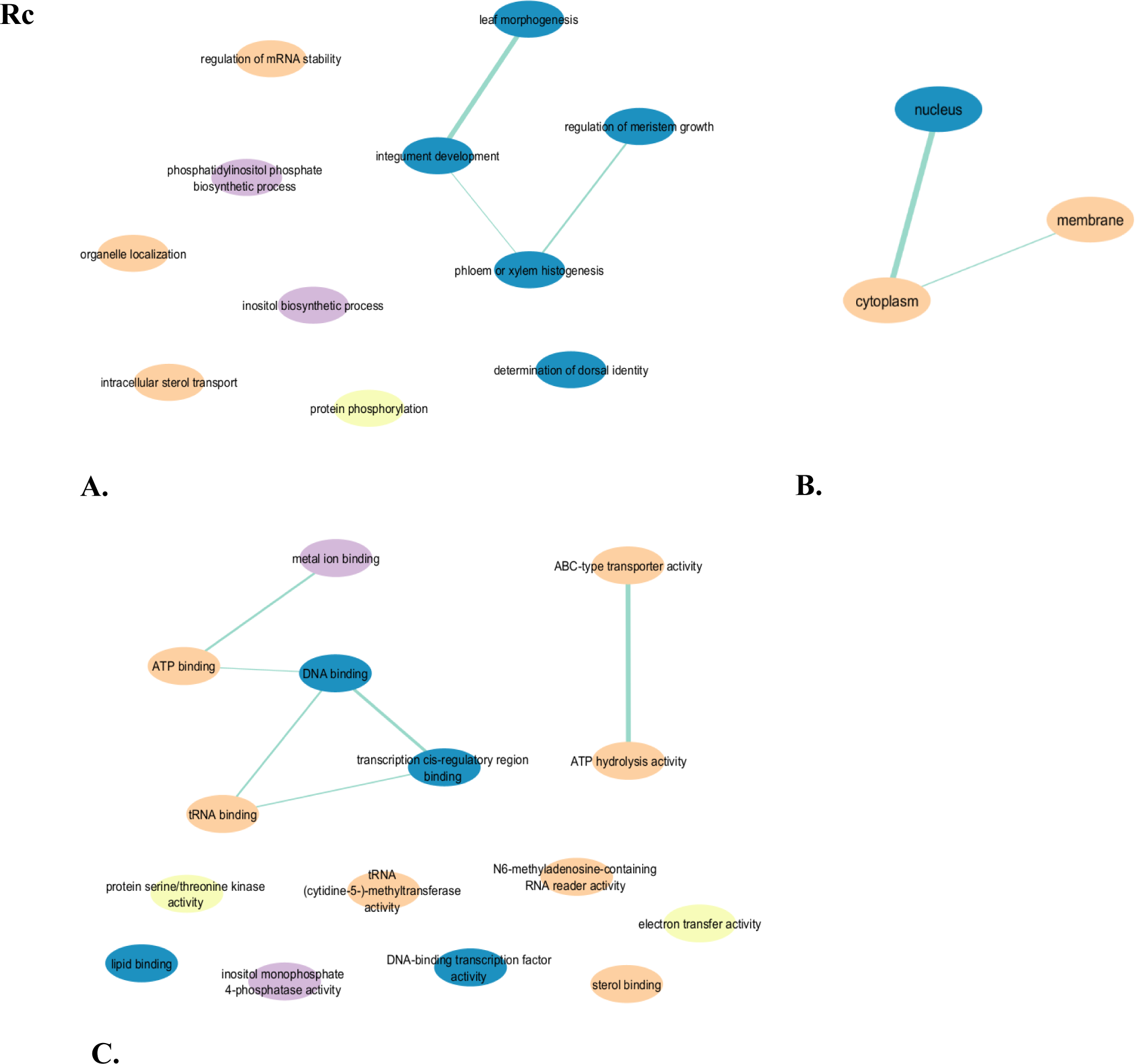
GO analysis of potential miRNA targets of the renewed seeds sample (Rc) for biological, cellular and molecular functions: **A.** biological functions, **B.** cellular functions, **C.** molecular functions.

**Figure 14.**
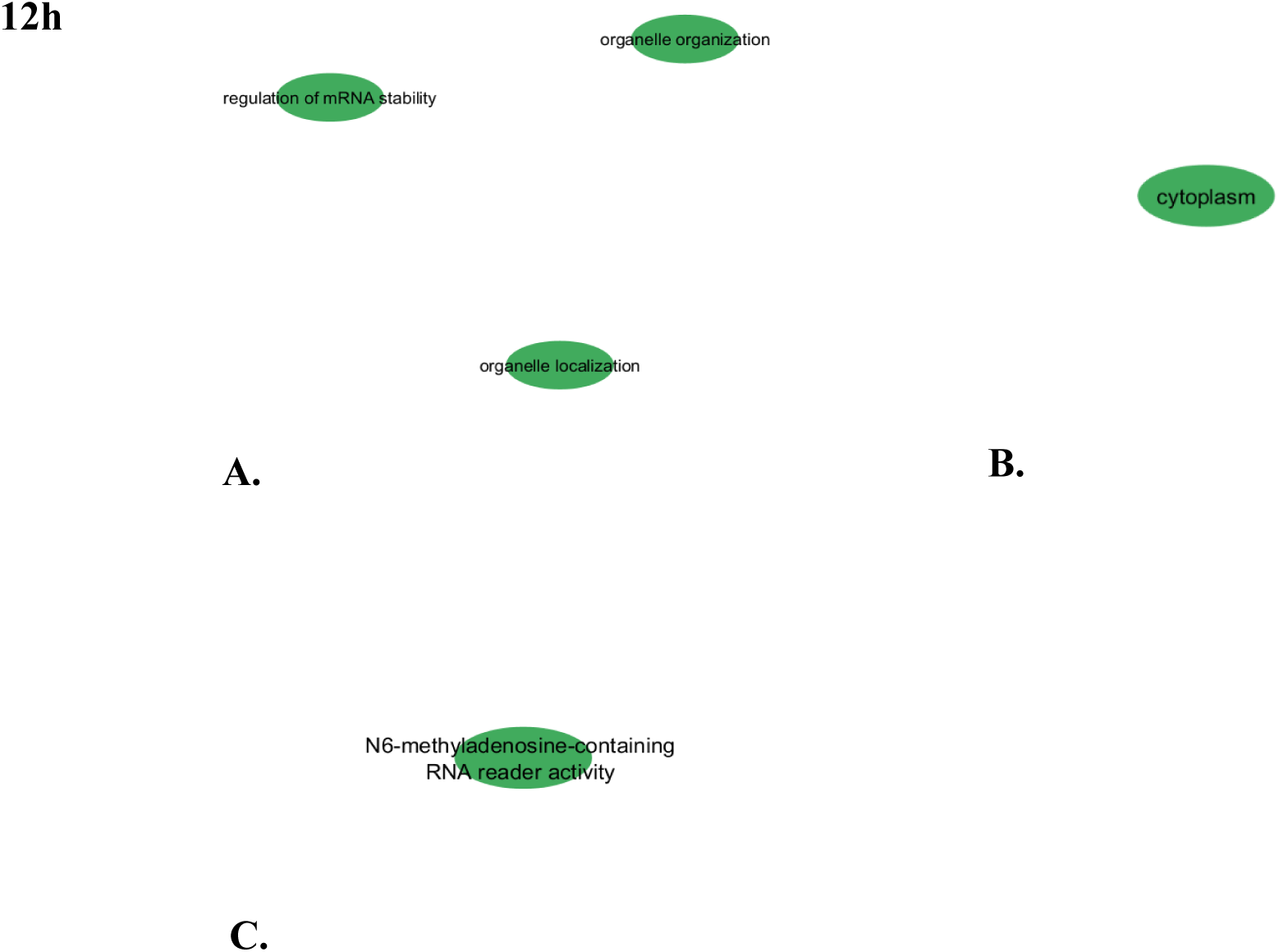
GO analysis of potential miRNA targets of the sample 12 hours after imbibition (12h) for biological, cellular and molecular functions: **A.** biological functions, **B.** cellular functions, **C.** molecular functions.

**Figure 15.**
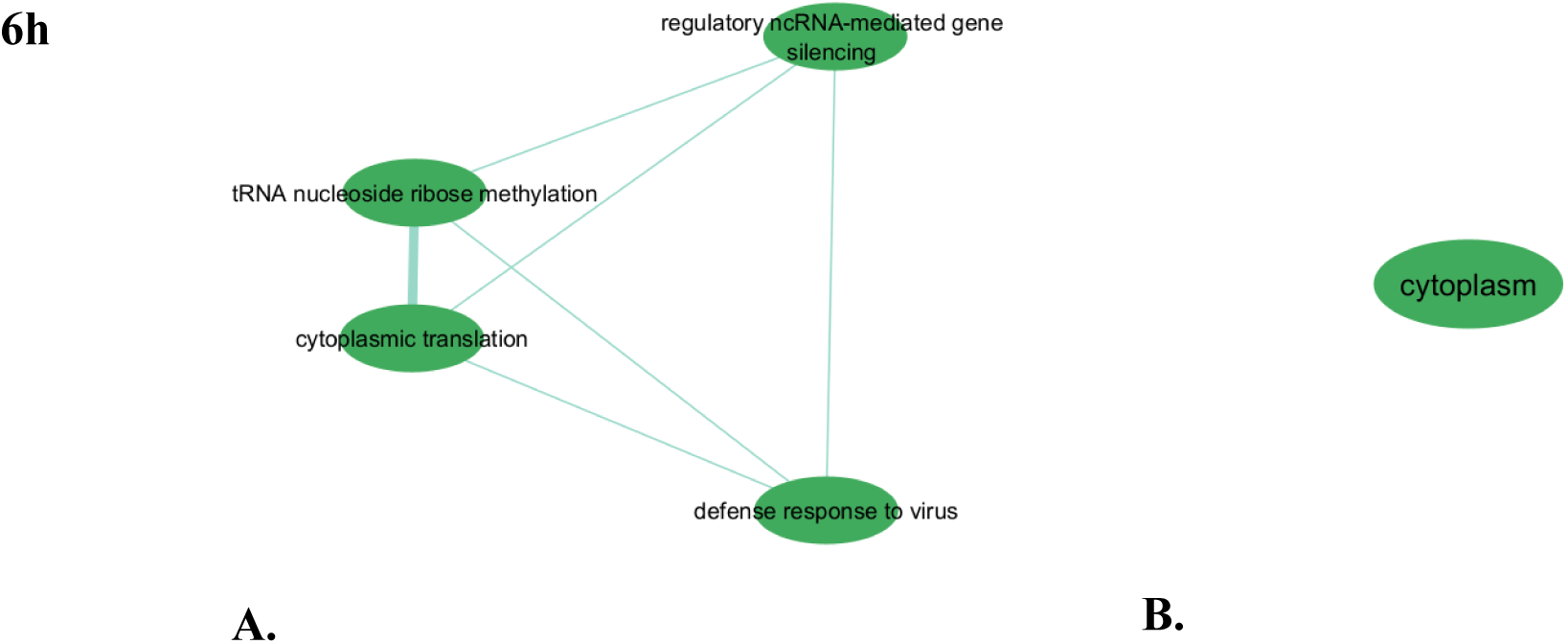
GO analysis of potential miRNA targets of the sample 6 hours after imbibition (6h) for biological, cellular and molecular functions: **A.** biological functions, **B.** cellular functions.

## Discussion

Small non-coding RNAs play an important role in germination and plant development. Until now, the process of germinating long-term stored seeds and how the miRNome changes during germination has not been studied. In this study, unique plant material was used, representing samples from a single long-term stored batch, which ensures that there were no differences due to the effects of environmental conditions. During long-term storage, one of the samples was unsealed and its moisture content increased, resulting in a drastic loss of viability. It was therefore possible to study the effect of seed vigor on the germination process by using naturally aged seeds with different levels of viability.

Loss of seed viability is associated with changes at the macromolecular level. These changes are most likely caused by reactive oxygen species and may interfere with the germination process [62]. The damage occurs to proteins as well as to DNA and RNA. RNA molecules are most susceptible to degradation during seed aging, and as viability declines, both the total amount of RNA and its integrity decrease [63–65]. However, the degradation of all RNA fractions is not uniform [66]. In particular, miRNA is the only fraction that does not degrade with seed aging [47]. This is due to the fact that in cells, miRNAs are incorporated into the RISC complex and are physically covered by it [67]. According to the EMBL-EBI Expression Atlas, the highest expression of RISC complexes has been associated with barley seed development and germination [67]. Therefore, it can be assumed that the starting point, i.e. the miRNA content, was homogenous for long-term stored seed samples tested here, and the differences that appeared during imbibition were related to aging and different germability.

The germination process is regulated by a complex network of regulatory mechanisms, with microRNAs (miRNAs) playing a pivotal role [68]. Our findings revealed distinct miRNA expression patterns in seed samples with varying levels of viability during the initial 24 hours of germination. The differential expression observed in these samples highlights the dynamic nature of miRNA-mediated regulation in response to seed viability and germination time. A total of 62 known microRNAs (miRNAs), belonging to 11 distinct families, and 234 novel miRNAs were identified in the aforementioned experiment. Regardless of viability and germination stage, 20 of these miRNAs were present in every sample tested. The known miRNAs included miR156, miR159, miR166, miR168, miR171, miR5048 and miR5051. Furthermore, as described by Puchta et al. [47], these miRNA families were also present in the dry seeds of the tested samples. The miR156, miR159, miR166, miR168 and miR171 families are highly conserved among land plants and performs various critical functions. They were identified in both monocotyledonous and dicotyledonous plants [69]. In general, all these miRNA families tended to be expressed at high levels, suggesting their involvement in important metabolic processes necessary for normal development and function.

miR156 is highly conserved in plants. Its target is the plant-specific transcription factor *Squamosa Promoter Biding Protein-like* (*SPL*) [70, 71]. The miR156-*SPL*s module regulates several age-dependent developmental processes and plays an important role in response to biotic and abiotic stimuli. Very high miR156 expression was also observed previously during barley seed development and germination [37, 72]. In total, eight different miR156 isoforms were observed during barley seed development. However, miR156a was significantly the most highly expressed [72]. In dry barley seeds with different levels of viability after long-term storage, only the miR156a isoform was identified, and its expression was comparable in all samples and was the highest of all the miRNAs identified [47]. Here, during germination of the aforementioned barley seed samples, miR156a was present only in fresh seeds after 6 hours of imbibition (Rc6). In rice, osa-miR156a knockout mutants had significantly slower germination, but no effect on shoot architecture was observed [73]. In *Arabidopsis*, the miR156a isoform, together with miR156c, is thought to play a dominant role in the family in response to salt and drought stress [74]. Fard et al. reported the involvement of miR156a during drought stress in barley, where it regulates the expression of genes involved in the dehydration response [75]. The high number of miR156a transcripts in dry seeds and in the early germination stage of the Rc sample may indicate its role in regulating the initiation and proper course of germination at a very early stage. Another isoform, miR156b, was predominant in seeds with low viability at the same time point (Lv6). Other isoforms except miR156a were also present in this sample at all early stages of germination. miR156f had high expression at Lv24, Hv12 and Rc6. In rice, the target of osa-miR156f is *OsSPL7*, and the miR156f/*OsSPL7* module regulates plant architecture. A mutant with up-regulated miR156f and down-regulated *OsSPL7* expression was identified. It had increased shoot number and reduced plant height [76]. Furthermore, the expression of tillering regulators such as *Teosinte Branched 1* (*TB1*) *and Lax Panicle 1* (*LAX1*) was downregulated by osa-miR156f [77]. This may therefore suggest that high expression of miR156f at early stages of germination determines normal coleoptile growth and prevents premature tillering. Its delayed expression in the low viabile seed sample may be due to the prolongation of germination stages [78]. And its effect may be an increased proportion of abnormal seedlings obtained from low viable seeds [79, 80].

miR159 is also an evolutionarily conserved family of miRNA present in most land plants, even described by some researchers as ancient [81]. The sequence of this canonical miR159 appears to remained constant over hundreds of millions of years, as very similar miR159 isoforms are present in most land plants [52]. It targets GAMYB or GAMYB-like transcription factors belonging to the MYB family, which are important gibberellin (GA)-induced regulatory proteins critical for grain development and germination and involved in anther formation [82–85]. GAMYB is not involved in GA signaling in vegetative tissues. Its expression results in inhibition of growth and development. Thus, miR159’s main function is to strongly silence GAMYB to allow normal growth [42]. In the early stages of barley seed development, mir159a was expressed. However, the level of expression was rather low [72]. miR159a was present in all dry barley seed samples but did not discriminate between samples with variable viability [47]. In contrast, during germination of the aforementioned seed samples, the presence of up to 9 isoforms of miR159 was detected. In the early stages of barley seed germination, the total expression of the miR159 family was one of the highest recorded. Like miR156, different isoform expression patterns were observed in different seed samples, however all had the same target. It is worth noting that the expression of miR159 increased during the first 12 hours of imbibition. Afterwards, a decrease was observed. It has already been reported that mir159 expression gradually increases as barley germinates and seedlings develop [37]. The above results indicate that miR159 levels fluctuate, which was also previously observed during barley development [86]. The initial fluctuation and subsequent increase in miR159 expression may be related to the regulation of starch reserve activation in the grain during germination, since in barley aleurone GAMYB positively transduces GA signaling to activate α-amylase and other hydrolytic enzymes expression and PCD in aleurone [87, 88]. Noteworthily, in rice, mature miR159 is absent in seeds despite being present throughout the plant [42]. In *Arabidopsis,* miR159 is upregulated in seeds during germination following the stress hormone ABA and drought [41]. In maize, wheat, and barley, miR159 also accumulates to higher levels in response to drought [24]. These findings may indicate that increasing miR159 levels may result in increased stress tolerance. This could be an additional explanation for the high level of miR159 expression that was observed in the very early stages of germination of the barley seed.

miR168 was highly expressed in the early stages of barley seed germination. In higher plants, this is another highly conserved family [89]. This family was also present in dry barley seeds. However, the number of transcripts was significantly lower [47]. The presence of mir168 during seed development and germination has also been demonstrated in other studies [37, 72]. Uniquely, miR168 is one of three miRNAs that control its own expression and the expression of other miRNAs. It targets the AGO1 protein involved in post-transcriptional gene silencing. Specifically, miR168 regulates the function of all miRNAs by modulating AGO1 levels and consequently RISC complex activity [90, 91]. It has been shown that AGO1 homeostasis, which is critical for miRNA-mediated post-transcriptional gene silencing, is maintained by a complex mechanism involving miR168-directed cleavage of AGO1 transcripts and miR168 stabilization through AGO1 association [92]. During imbibition, the seed activates a series of genes that are responsible for the restoration of cellular integrity, the repair of mitochondrial and DNA damage, as well as the initiation of respiration and metabolic activity, the use of reserve compounds [93]. Transcriptional, post-transcriptional and post-translational regulatory mechanisms, including miRNAs, are involved in this process [93]. This study showed that the composition of the microtranscriptome is significantly enriched during the germination of barley seeds as compared to dry seeds. Thus, during early stages of barley seed germination, the high level of miR168 expression responds to the need to regulate the activity of other miRNAs. Stable mature molecules from both pre-miRNA arms were found in dry seeds for miR168a-5p/168a-3p [47]. Similar results were previously obtained in rice, maize and barley [86]. Notably, in here, miR168a was only present during germination of high viable (Hv) seeds after 12 and 24 hours of imbibition. It was absent either in poorly viable seeds (Lv6-24) or during the first 6 hours of germination (Hv6). A total of 12 miR168 isoforms were identified in the early stages of barley seed germination. This makes this family the most diverse of all detected miRNAs. Monocotyledons have a higher diversity of miR168 than dicotyledons, as shown by Jagtap et al.[94]. Simultaneously, there is also a higher number of AGO1 members [94]. Notably, miR168 expression was almost absent in the Lv6 sample. Considering that miR168 regulates AGO1 expression, lack of expression of this miRNA may indicate high levels of AGO1 in this seed sample. As discussed earlier, modulating AGO1 levels affects the activity of the RISC complex. Thus, the high total miRNA expression in the Lv6 sample and possibly high AGO1 expression may be an indication of much more intense post-transcriptional gene regulation during the early germination phase of low viable seeds. However, further RNAseq analysis during early germination of seeds with different viability is needed to verify this hypothesis.

miR444 family is specific to monocotyledonous plants. It has been identified in rice, wheat, barley, maize, sorghum, tussock and sugarcane [95]. It has also been reported that miR444 targets genes encoding MADS-box family transcription factors [95, 96]. In rice, miR444 targets five homologous MADS-box transcription factors (MADS23/25/27/57/61) involved in the regulation of brassinosteroids (BR) biosynthesis. The TFs negatively regulate BR biosynthesis by binding directly to the *OsBRD1* promoter. Thus, miR444-MADS-box-*OsBRD1* controls BR homeostasis as a key signaling cascade [97]. After 6 hours of imbibition, miR444 isoforms were present only in the low viable seed sample (Lv). Expression was absent in the other two samples, although miR444b was present in dry seeds [47]. Thus, as vigor decreased, upregulation was characteristic. Similarly, maize zma-miR444 was upregulated as seed vigor decreased. Negative regulation of miR444 and its target gene *AP2-ERF* was found to be associated with maize seed storage via the ethylene pathway [98].

Deng et. al (2016) explained the high expression of miRNAs by their role in barley growth and development. They observed a difference in the number of miRNAs identified: some were only a few, while others were thousands [99]. The conclusions of the researchers are in line with the observations from the analysis of the results presented here. Differences in the expression levels of miRNA families may be due to the activity of different physiological and biochemical processes in the seeds at any given time of imbibition [100]. Differences in the expression pattern of individual miRNAs may indicate which pathways are activated during germination. The same expression trend was observed in the results obtained by qRT-PCR and NGS sequencing analyses. In the study described herein, a considerable number of novel microRNAs were identified - specifically, 234. Prior studies on barley seed development and germination have likewise identified numerous novel microRNAs. A common feature of the aforementioned novel microRNAs is their low level of expression [37, 101]. This is in agreement with the results of other authors, although results may differ due to the sensitivity of the methods [102].

Seed imbibition triggers many biochemical and cellular processes associated with germination, including reactivation of cellular respiration and metabolism, DNA repair, transcription and translation of new mRNAs, which coincides with the functions of miRNA target genes with significant expression differences observed during the study. The results obtained here indicate the initiation of processes necessary for the resumption of metabolism and the involvement of miRNAs in their regulation. Functional analysis of genes regulated by known miRNAs revealed that, regardless of their viability level, they are mainly involved in the regulation of protein modification and especially in phosphorylation. Protein phosphorylation is a reversible post-translational modification. A kinase adds a phosphate group to a protein. In signal transduction and enzyme regulation, this process is fundamental. Phosphorylation activates or deactivates proteins involved in signaling pathways necessary for response to environmental factors and hormonal signals during germination. Phosphorylation also alters enzyme activity. Thus, it controls metabolic pathways required for energy production and biosynthesis [103]. Although the underlying mechanisms remain poorly understood, it has been reported that the association of different classes of miRNAs with AGO proteins during RISC loading appears to be phosphorylation dependent [104]. The potential to integrate miRNA biogenesis into cellular responses to stress and development exists through transcriptional regulation of individual MIR loci through binding of phosphoproteins for example hyponastic leaves 1 (HYL1) in the DCL1 complex [105]. Thus, miRNAs that regulate phosphorylation can regulate the biogenesis, activity and efficiency of subsequent miRNAs during seed germination. Also in all seed samples, miRNA-regulated genes were involved in phosphorus metabolism and phosphate-containing compound metabolism. Phosphorus, which plays a critical role in energy transfer, signal transduction, and the synthesis of nucleic acids and membranes, is an essential element for plant growth and development. During seed germination, phosphorus and phosphate metabolic processes are particularly important for the many biochemical and physiological changes required to transition from dormancy to active growth including energy transfer, nucleic acids synthesis, phospholipids biosynthesis and signal transduction. During the germination process, adenosine triphosphate (ATP) is produced through cellular respiration and utilized as a fuel for various metabolic activities. The hydrolysis of ATP to adenosine diphosphate (ADP) releases energy that drives the biochemical reactions necessary for seedling growth [106–108]. Phosphorus is also an indispensable component of the backbone of both DNA and RNA molecules. During the germination process, the synthesis of new DNA and RNA is vital for cell division and protein synthesis. A sufficient supply of phosphorus is essential for the efficient production of nucleic acids, which in turn supports the growth and development of the seedling [109]. The synthesis of nucleotides depends on phosphorus. They are not only essential for genetic material but also serve as energy carriers (e.g., ATP, GTP) and cofactors in enzymatic reactions [110]. Moreover, phosphorus is a pivotal element in the head groups of phospholipids, playing a crucial role in maintaining membrane integrity and functionality. During the germination process, the synthesis of new membranes is of critical importance for the expansion of cells and the formation of new cellular structures [111]. It is notable that the regulatory function of microRNAs on the genes involved in organonitrogen compound metabolism was only observed in seeds with low viability after six hours of imbibition, as evidenced by the results of the Gene Ontology (GO) analysis. The metabolic processes involved in the synthesis, transformation, and degradation of nitrogen-containing organic molecules, collectively referred to as “organonitrogen compounds,” represent a fundamental aspect of biochemical activity within living systems. These processes play an essential role during the germination of seeds, as nitrogen serves as a fundamental component in the synthesis of amino acids, nucleotides, and other vital cellular constituents. Amino acids serve as the fundamental units for the synthesis of proteins. During germination, the synthesis of new proteins is vital for cellular functions, including enzymatic activity, structural integrity, and signal transduction. The assimilation of nitrogen into amino acids, including glutamine, glutamate, and asparagine, is a critical step in protein biosynthesis [111, 112]. One of the primary factors affecting seed longevity is the deterioration of proteins that serve a multitude of functions in seeds, including structural and enzymatic roles. This damage can result from a number of different mechanisms, including oxidation, deamination, glycation, and fragmentation. Protein damage directly affects the ability of seeds to germinate. Enzymes involved in the germination process may be inactive and cellular structures damaged, leading to reduced seed vitality [113–115]. This suggests that miRNA may play a key role in regulating the mechanisms of repairing protein damage caused by seed aging.

The translational capacity of stored mRNA is a determinant of seed germination during storage. Dry seeds are in translation quiescence. However, it has been reported that approximately 50% of seed-stored mRNA is associated with ribosomes and is translationally regulated during imbibition.. In Arabidopsis age-resistant genotypes, high expression of genes responsible for protein elongation, mobility of the tRNA molecules was observed. These genes regulate the translation process by facilitating it for ribosome-associated mRNAs during the early stage of seed germination [100]. During the study, genes related to nucleic acid binding processes were observed, while degradome sequencing identified a gene related to the ribosomal protein l10(RPL10), which belongs to the large subunit of the ribosome [116]. The number of proteins that can be synthesized depends on the number of ribosomes, so ribosome biosynthesis is a process that determines the development of an organism. During the first 24-48 h of imbibition, there is a high demand for ribosome production and protein synthesis, which is related to the penetration of the embryonic axis and the surrounding structures [117]. The first phase ends with a plateau phase (phase II) involving the activation of a number of metabolic processes that facilitate energy production and reserve mobilization. During this process, ribosomal protein gene expression and ribosomal activity increase dramatically, facilitating the new synthesis of proteins important for seed germination [118]. In the Arabidopsis study, the researchers observed that miRNAs were prepared during seed maturation for subsequent fate during germination. They show that ∼50% of seed-stored mRNA is associated with ribosomes, mainly monosomes [119], our results also indicated the involvement of miRNA in ribosomes development.

### Conclusions

The results presented here clearly demonstrate the involvement of miRNAs in the germination process. The constitutive miRNA families maintain a constant level of expression at all stages of germination and are essential for normal metabolic processes in the plant. Among the identified miRNAs, isomiRs with tissue-specific expression patterns were observed, suggesting the occurrence of distinct metabolic processes in seeds of different viability levels at the same time point during imbibition. A decline in total miRNA expression was observed in all seed samples after 24 hours of imbibition. This suggests that the restoration of normal cellular metabolism in quiescent and desiccated seeds may be associated with increased transcriptional activity and the need for transcriptional regulation by miRNAs. Seeds that were found to retain high viability after long-term storage exhibited a nearly twofold increase in total miRNA expression compared to a sample of freshly regenerated seeds. However, a decline in isoforms diversity was observed. This suggests that the process of germination following long-term storage requires a significantly elevated level of transcription and miRNA associated regulation. This may be due to the necessity to initiate repair mechanisms for macromolecules or cellular ultrastructure.

Analysis of the function of the target genes of the identified miRNAs revealed their association with translation processes and ribosome function, which may be crucial for maintaining viability and initiating the germination process. Although the mechanism of miRNA involvement in the germination process of long-term stored seeds remains to be investigated, the results provide new insights into the molecular mechanism underlying the regulation of the germination process.

## Supporting information

Table 1. List of sequences primers for RT-qPCR

Table 2. An average level of expression of know miRNA identified in the tested seed sample

Table 3. An average level of expression of novel miRNA identified in the tested seed sample

## List of abbreviations

miRNA: microRNA
sRNA: small RNA
Lv: seeds with low viability
Rc: renewed seeds
Hv: seeds with high viability
NGS: next generation sequencing
RT-qPCR: quantitative reverse transcription-PCR
OPPP: oxidative pentose phosphate pathway
TCA: tricarboxylic acid
siRNA: interfering RNAs
RPM: reads per million
GO: gene onthology
RT-qpcr: reverse transcription qPCR
TB1: Teosinle Branched 1
LAX1: Lax Panicle 1
BR: brassinosteroids

## Ethics approval and consent to participate

Experimental research and field studies on cultivated plants, including the collection of plant material, are carried out in accordance with all applicable institutional, national, and international guidelines and legislation throughout the course of this study.

## Consent for publication

Not applicable.

## Authors’ contributions

Conceptualization, M.P.J.; M.B. and J.G.; methodology, M.P.J., J.G.; formal analysis, M.P.J.; P.B. and J.G.; data curation, M.P.J.; writing—original draft preparation, M.P.J. and P.B.; writing—review and editing, M.B. and J.G.; visualization, M.P.J.; supervision, M.B.; funding acquisition, M.P.J. All authors have read and agreed to the published version of the manuscript.

